# A mathematical model of osteocyte network control of bone mechanical adaptation

**DOI:** 10.1101/2025.10.10.681578

**Authors:** Adel Mehrpooya, Vivien J Challis, Pascal R Buenzli

## Abstract

The osteocyte network embedded in bone tissues plays a central role in the control of bone adaptation to mechanical loads and micro-damage repair. However, much remains to be understood about the precise mechanisms by which the osteocyte network regulates bone formation and bone resorption based on the propagation of biochemical signals emitted in response to mechanical stimulus. In this work, we propose a one-dimensional computational model of bone mechanical adaptation based on the propagation of signalling molecules through a dynamic osteocyte network. The osteocyte network is extended during bone formation, and reduced during bone resorption, which affects the generation and propagation of the signalling molecules to the bone surface. We explore how this osteocyte-based model of bone mechanosensation and mechanoresponse gives rise to effective Wolff’s laws, in which overloaded bone is consolidated and underloaded bone is removed. We find that the discrete addition and removal of osteocytes significantly affects signalling molecules propagating to the bone surface and leads to new bone adaptation behaviours compared to earlier models, including partial bone recovery following an unloading and reloading cycle, and the emergence of a minimum threshold of mechanical stress below which all bone is resorbed. While many extensions of this mathematical model are possible, it provides a first illustration of how the osteocyte network could act as a dynamic embedded control network for bone adaptation to mechanical loads.

## 1 Introduction

Bone is an adaptive biological tissue that fulfills important mechanical functions (Martin, Burr & Sharkey 1998; Cowin 2001). To be light and strong, bone optimises its shape in response to mechanical loading, such that regions experiencing high loads are consolidated and regions experiencing low loads are removed (Bertram & Swartz 1991; Turner 1999; Hernandez, Carter & Beaupré 2000; Carter & Beaupré 2001; Frost 2003; Skerry 2008; Rosa et al. 2015). Bone tissues also undergo continual self-repair to prevent the accumulation of fatigue micro-cracks (Martin 2003). Old, damaged tissue is periodically replaced with new tissue by a process called bone remodelling. Bone adaptation and bone remodelling are fundamental biological processes that allow our bones to be mechanically under-engineered without risking spontaneous fracture (Martin 2003).

There is increasing evidence that the osteocyte network embedded in bone tissue is central to the orchestration and control of these adaptative processes (Mullender et al. 1996; Qiu et al. 2002a,b; Vashishth et al. 2000; Chen et al. 2010; Tiede-Lewis & Dallas 2019).

Osteocytes are mechanosensitive cells that live in bone tissue within a fluid-filled porous network called the lacunocanalicular network (LCN) (Bonewald 2011; Franz-Odendaal, Hall & Witten 2006; Mader et al. 2013; Buenzli & Sims 2015; Kollmannsberger et al. 2017; Weinkamer, Kollmannsberger & Fratzl 2019). The presence of osteocytes within hard bone tissue enables these cells to identify regions needing consolidation, removal, or repair, and to emit signals in response to mechanical stimulus to induce bone formation and bone resorption.

Mechanical loads applied to bone tissues induce fluid flow within the LCN that generates shear stress on the osteocyte cell bodies, triggering responses by osteocytes through mechanobiological pathways. This so-called fluid-flow hypothesis (Weinbaum, Cowin & Zeng 1994; Turner & Forwood 1995; Knothe Tate et al. 2000; van Tol et al. 2020a,b; Fu & Yang 2024) is believed to be a primary mechanism for bone mechano-sensation, although osteocytes may also respond to a variety of mechanical variables such as hydrostatic pressure, strain, and strain rate (Meakin, Price & Lanyon 2014; Wang, Fu & Yang 2025). Osteocytes respond to these mechanical stimulations by emitting signalling molecules, such as sclerostin, RANKL and other paracrine factors (Turner & Forwood 1995; Robling, Castillo & Turner 2006; Robling & Turner 2009; Ryser Nigam & Komarova; Xiong et al. 2011; Nakashima et al. 2011; Moriishi et al. 2012; Lau et al. 2013). Gap junctions between osteocyte cytoplasmic processes also provide cell-cell communication of intracellular ions and cytokines triggered by mechanical stimulation, such as calcium and cyclic AMP (Robling & Turner 2009; Loiselle, Jiang, Donahue 2013; Lewis et al. 2017; Palumbo & Ferretti 2021). While the LCN is often assumed to influence mechano-sensation via the fluid-flow hypothesis, the osteocyte network of direct cell–cell contact provides an independent means of intercellular communication by which mechano-response signals may propagate. When these signals reach the bone surface, they promote the generation of osteoblasts (bone-forming cells) or osteoclasts (bone-resorbing cells) (Marotti 2000; Franz-Odendaal, Hall & Witten 2006; Bonewald 2011; Klein-Nulend & Bacabac 2012; Manolagas & Parfitt 2013; Palumbo & Ferretti 2021; Ferretti & Palumbo 2021).

Many computational models of bone adapation to mechanical loading are based on Wolff’s law (Huiskes et al. 1987; Mullender et al. 1995; Carter & Beaupré 2001; Jang & Kim 2010; Schulte et al. 2013; Wang et al. 2020, 2023). These models are successfully used in orthodontics and orthopaedics to predict the stability of screws and implants (Chou, Jagodnik & Müftü 2008; Ferguson et al. 2022; Wan et al. 2022) and to simulate changes in the shape and microstructure of bone subjected to physical loads (Mullender et al. 1995; Huiskes et al. 2000; Jang & Kim 2010; Adachi et al. 2010; Farhoudi et al. 2017; Fallahnezhad et al. 2017). Such models may be coupled with pharmacokinetic models to simulate osteoporosis therapies targeting osteocytes (Joerg et al. 2022). These computational models usually assume a fixed reference mechanical state, called the mechanical setpoint. Few studies investigate the generation of osteocytes (Buenzli 2015, 2016; Taylor-King et al. 2020) and specific links between the cellular network of osteocytes and bone’s ability to sense and respond to mechanics (Lerebours & Buenzli 2016; Pauchard & Buenzli 2025). While some works include osteocyte signalling in bone adaptation models (Mullender et al. 1995; van Oers et al. 2008; Adachi et al. 2010; Adachi & Kameo 2018; Prasad & Goyal 2019), they do not account for the fact that the osteocyte network itself is dynamic due to bone formation and resorption, which influences how signals propagate through the network to the bone surface. How the osteocyte network acts as a regulatory control network for bone adaptation while itself being affected by adaptation is an important question that mathematical modelling can help answer.

In this work, we propose a one-dimensional computational model of bone mechanical adaptation based on the propagation of signalling molecules through a dynamic osteocyte network. The molecules are emitted by osteocytes in response to mechanical stimulation. They propagate to the bone surface, where they induce the formation of osteoblasts (bone-forming cells) or osteoclasts (bone-resorbing cells). The osteocyte network is extended during bone formation, and reduced during bone resorption. These processes affect the generation and propagation of the signalling molecules. We explore how this osteocyte-based model of bone mechanosensation and mechanoresponse gives rise to effective Wolff’s laws, in which overloaded bone is consolidated and underloaded bone is removed. The aim of our study is to use this model of signal propagation to analyse qualitatively (i) how osteocyte regulation of bone formation and resorption affects the extent and propagation properties of the osteocyte network itself; (ii) how mechanical adaptation depends on the type of mechanical stimulus assumed to trigger osteocyte responses; and (iii) how osteoblasts and osteoclasts may respond to signals emanating from the network’s edges at the bone surfaces. Our model builds on our previous work of signal propagation through (static) one-dimensional spatial networks using stochastic and deterministic random walks and their continuum limit (Mehrpooya, Challis & Buenzli 2024). While the model we propose represents a simplified view of bone mechanical adaptation, it reveals useful insights into processes underlying bone mechanosensation and mechanoresponse that provide a first illustration of how the osteocyte network acts as a regulatory embedded control network.

## 2 Mathematical Model

### 2.1 Signal propagation in the osteocyte network

In this section we present a model of the propagation of signalling molecules within a one-dimensional, dynamic osteocyte network embedded in bone tissue. This model is based on the random walk model proposed by Mehrpooya, Challis & Buenzli (2024), however we restrict the current work to a deterministic model and do not consider stochasticity. The network can be considered to represent the osteocyte network in a one-dimensional slice *b*_−_(*t*) ⩽ *x* ⩽ *b*_+_(*t*) through the cortical wall of a mouse tibia, where the position of the left bone boundary *b*_−_ (*t*) and the right bone boundary *b*_+_(*t*) can vary with time *t*, as shown schematically in Figure 1.

**Figure 1.**
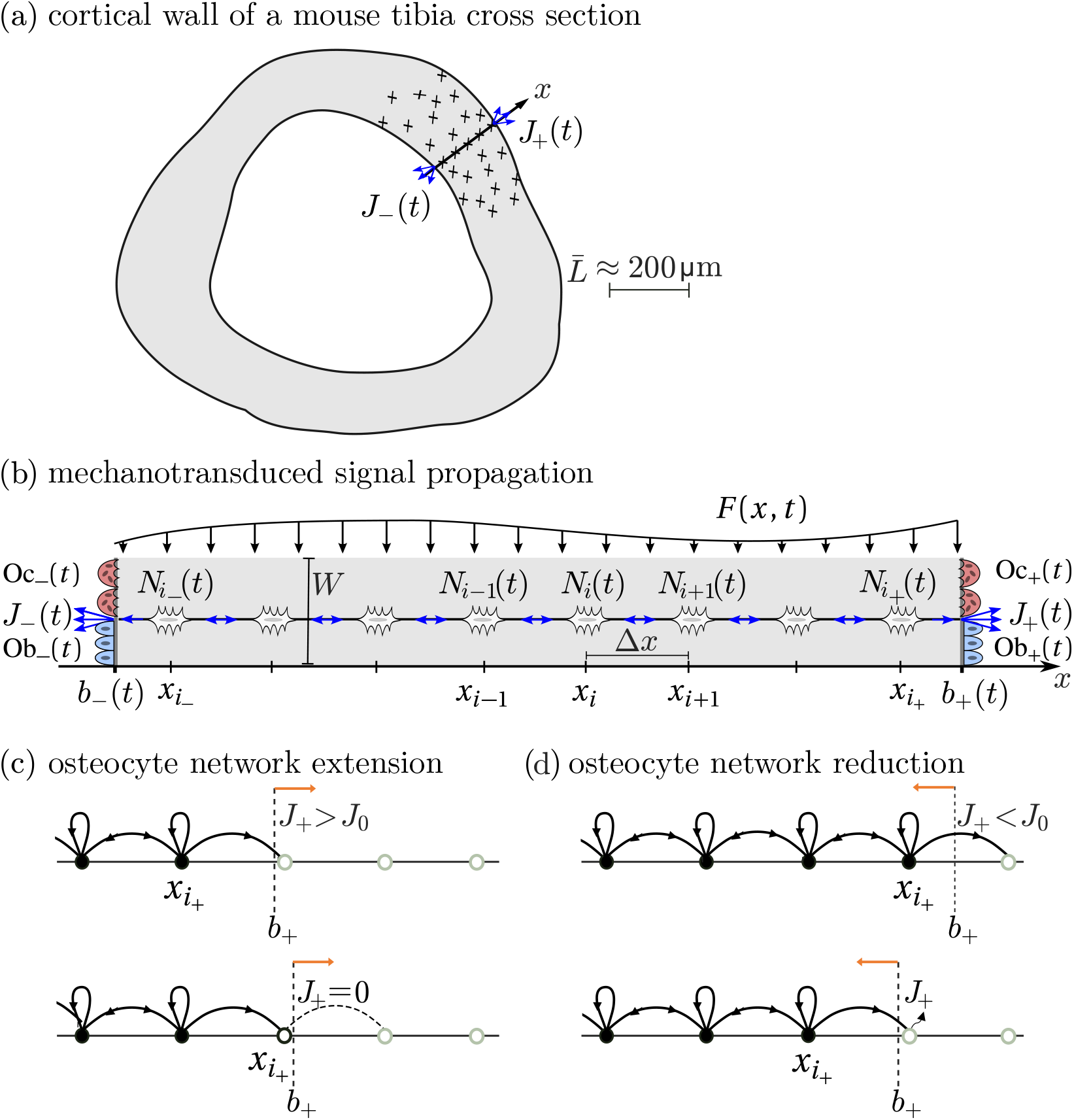
(a) Two-dimensional cross-section of a mouse tibia cortical wall (grey) showing the *x*-axis as a slice across the cortical wall. Osteocytes (black pluses) along this axis emit and propagate signalling molecules to the bone surface, where they induce signalling fluxes *J*_−_ (*t*) and *J*_+_(*t*). (b) Osteocytes are placed at equal distance Δ*x* from each other along *x*. The *i* ^th^ osteocyte has a number of signalling molecules *N*_*i*_ (*t*). The slice is assumed to be stimulated mechanically by a space and time-dependent loading force *F*(*x, t*). Signalling molecules propagate between osteocytes until they reach the bone boundaries *b*_±_(*t*). The fluxes *J*_±_(*t*) of signalling molecules emanating from the bone surface induce the generation of osteoblasts Ob_±_(*t*) (bone-forming cells) or osteoclasts Oc_±_(*t*) (bone-resorbing cells). (c) Evolution of the osteocyte network during bone formation. Before the network extends (top), the flux is greater than the reference value, i.e., *J*_+_ *> J*_0_. When a new osteocyte is formed (bottom), it is initialised with zero signalling molecules, so *J*_+_ = 0. (d) Evolution of the osteocyte network during bone resorption. Before the network reduces (top), the flux is less than the reference value, i.e., *J*_+_ *< J*_0_. When an osteocyte is removed (bottom), the flux *J*_+_ includes molecules jumping right from the new right-most osteocytes and molecules that were previously in the resorbed osteocyte

Our focus is on the adaptation of bone via the time-evolution of bone boundaries regulated by the osteocyte network. The network is assumed to be represented by regularly spaced osteocytes with spacing *Δx*. Each osteocyte is labelled by a discrete index *i* = *i*_−_, *i*_−_ +1, …, *i*_+_ −1, *i*_+_ and is located at position *x*_*i*_ = *i*Δ*x* (Figure 1b). We denote by *i*_−_ the index of the left-most osteocyte at time *t*, and by *i*_+_ the index of the right-most osteocyte at time *t*. These indices depend on time due to the dynamic population of osteocytes, see Section 2.2.

We denote the quantity of signalling molecules positioned at osteocyte *i* at time *t* by *N*_*i*_ (*t*) (Figure 1b). Molecules are assumed to be generated by osteocytes in response to a mechanical stimulus. The rate of molecule production at osteocyte *i* is denoted by *R*(*x*_*i*_, *t, N*_*i*_) (number per unit time). Signalling molecules are also assumed to propagate to neighbouring osteocytes. We define 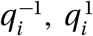 and 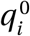 as the fractions of molecules that move left, right, or remain stationary at osteocyte *i*, respectively, during the time interval Δ*t*. These fractions are such that 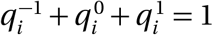. While longer-range jumps could be considered, representative signalling behaviour occurs when including only these nearest-neighbour jumps (Mehrpooya, Challis & Buenzli 2024). With the above notation, the evolution of the molecular occupancy is described by the update rule

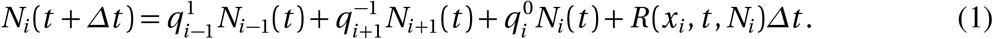

This update rule applies for *i*_−_ ⩽ *i* ⩽ *i*_+_ with the convention that 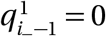 and 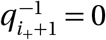, meaning that molecules can jump out of the network at the left and right bone boundaries, but not re-enter the network. This represents absorbing boundary conditions and the fact that signalling molecules coming out of the osteocyte network are processed by the micro-environment to induce bone formation or bone resorption. The flux of molecules coming out of the bone surface at the left and right boundaries of the network (number per unit time) is given by

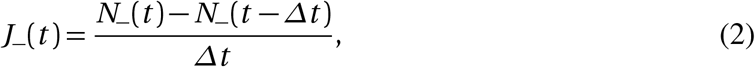

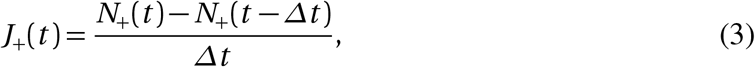

where *N*_−_(*t*) and *N*_+_(*t*) are the cumulative numbers of molecules that jump out of the network up until time *t* at the left and right boundary, respectively. These numbers include molecules jumping out of the network by Eq. (1), and molecules released into the microenvironment by resorption of the left-most or right-most osteocyte (see Figure 1d and Section 2.2).

Osteocytes may respond to a variety of mechanical stimulations Meakin, Price & Lanyon (2014); Wang, Fu & Yang (2025). We propose a mechanotransduction model in which the more mechanically stimulated osteocytes are, the more they produce signalling molecules. We consider two models for the molecule production rate *R*(*x*_*i*_, *t, N*_*i*_) in Eq. (1). The first assumes that osteocytes are stimulated by mechanical stress, such that the production of signalling molecules is

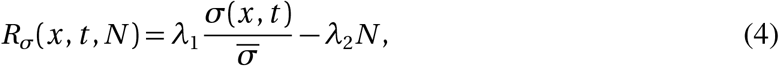

where *σ*(*x, t*) denotes the mechanical stress at position *x* and time *t*, 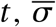 denotes a reference stress, and *λ*_1_ *>* 0 (day^−1^) and *λ*_2_ *>* 0 (day^−1^) are rates of molecule production and degradation, respectively. The stress is computed as force per unit area using

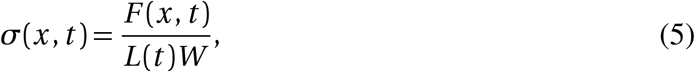

where *F*(*x, t*) is the applied mechanical force on the bone slice at position *x* and time *t, L*(*t*) = *b*_+_(*t*) − *b*_−_(*t*) is the cortical wall thickness, and *W* = 20µm is the assumed distance between osteocytes in the lateral direction (Qin et al. 2020) (see Figure 1b). The choice of *W* affects the scale of the appropriate force *F* to apply but is otherwise irrelevant.

Our second mechanotransduction model assumes that osteocytes are stimulated by the strain energy density, such that the production rate of signalling molecules is

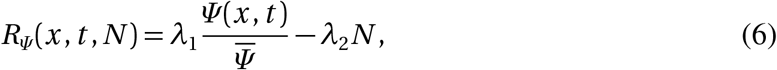

where *Ψ*(*x, t*) is the strain energy density and 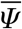 denotes a reference strain energy density. The strain energy density can be computed as

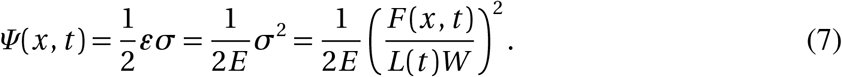

Here, *ε* is the strain and *E* is Young’s modulus. These quantities are related to the stress *σ* via Hooke’s law *σ* = *E ε*.

To determine reference values for the above mechanics-related parameters, we use data from the literature. These values are summarised in Table 1. We take the reference cortical wall width of a mouse tibia to be 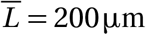, and assume that osteocytes are spaced at intervals of Δ*x* = 10µm (van Tol et al. 2020a; Mehrpooya, Challis & Buenzli 2024). The Young’s modulus of a mouse tibia is typically reported as ranging between 10 GPa and 15 GPa (Oliviero et al. 2021), and we take a value of *E* = 10 GPa. With this value of *E* and a reference strain value of 10^3^ *µε* (microstrains), we obtain a reference stress value of 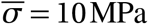 MPa, which is reasonable for a mouse tibia (van Tol et al. 2020a). From Eq. (5), the value of the force that gives a stress of 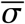 is therefore

**Table 1.**
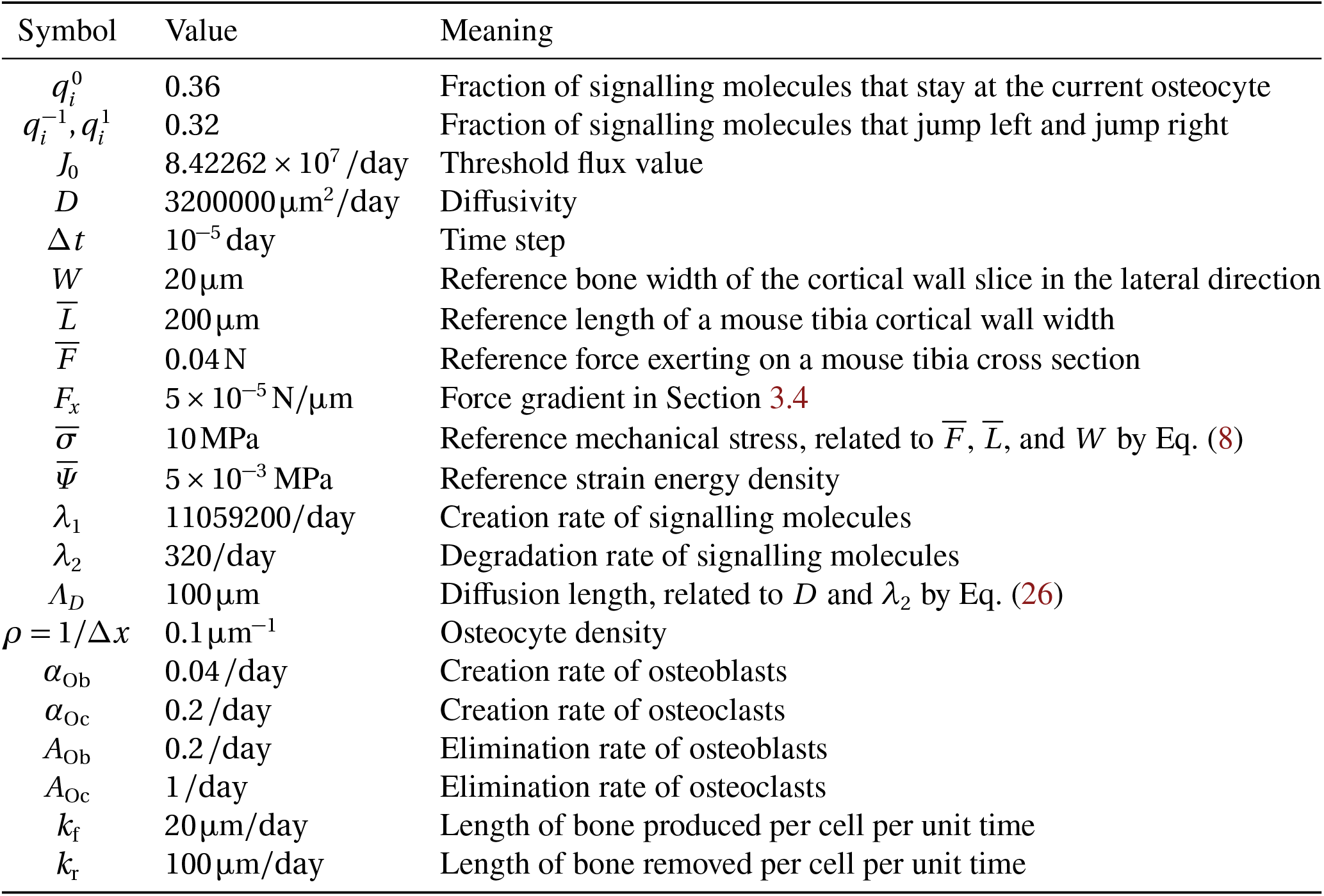
List of all parameter values of the discrete model with differential cell generation.

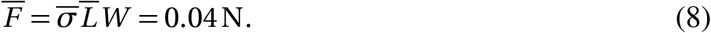

The same reference force applies in the case of Eq. (6), with 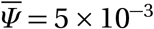 MPa computed from Eq. (7) using *E* and 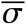.

It will be useful to consider the continuum limit of this discrete model of signalling molecule propagation. For this, we introduce the concentration *n*_*i*_ (*t*) of molecules at osteocyte *i* at time *t*, given by

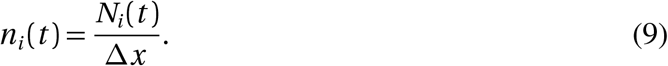

In Mehrpooya, Challis & Buenzli (2024), we showed that in the limit where Δ*t* and *Δx* approach zero, Eq. (1) converges to the following reaction–advection–diffusion equation:

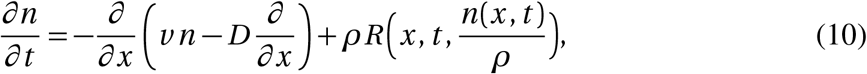

where *n*(*x, t*) represents the concentration of signalling molecules at position *x* and time *t*, such that *n*(*x*_*i*_, *t*) = *n*_*i*_ (*t*). In Eq. (10), *ρ* = 1/Δ*x* is the osteocyte density, *D* is the diffusivity and *v* is the advective velocity, obtained in the continuum limit as

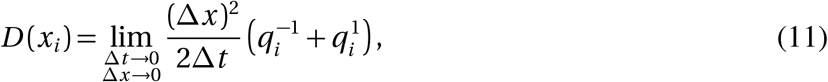

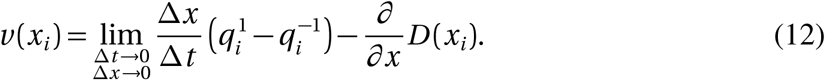

Here we will assume homogeneous and unbiased jumps, such that 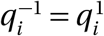 and 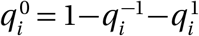 are independent of *i* for all *i*. In this case, *D* is constant and *v* = 0. Dirichlet boundary conditions

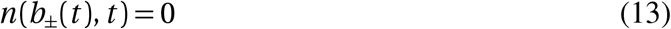

are imposed, which are equivalent to the absorbing boundary conditions described above for the discrete model by setting 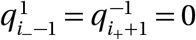.

### 2.2 Bone adaptation in response to osteocyte signals

In this section we describe our model for the adaptation of bone and of the osteocyte network in response to changes in mechanical loads. As shown schematically in Figure 1, we assume that the flux of signalling molecules at the boundaries of the network induces bone formation or bone resorption and corresponding evolutions of the bone boundaries.

According to our mechanotransduction model in Eqs (4) or (6), the production of signalling molecules is proportional to the mechanical stimulus, and results in a certain flux of molecules at the boundary. Therefore, a high flux of signalling molecules emanating from the bone surface should induce bone formation, while a small flux of signalling molecules should induce bone resorption. We assume that there is a reference flux value *J*_0_ at which the bone boundaries remain stationary. If the flux of signalling molecules at the bone surface exceeds *J*_0_, then bone is overloaded and osteoblasts are generated to form bone. If the flux of signalling molecules is less than *J*_0_, then bone is underloaded and osteoclasts are generated to resorb bone.

It is important to note here that *J*_0_ is not a traditional mechanical setpoint, in the sense that it is not a local bone material property that can directly be compared with the local mechanical state of bone matrix (Lerebours & Buenzli 2016). The reference flux *J*_0_ is associated with how osteoclasts and osteoblasts respond to mechanotransduced signals emanating from the bone surface, rather than with how osteocytes transduce local mechanical strains into biochemical signals. This distinction is absent in Wolff’s-type laws in which there is a direct relationship between local mechanical stimuli and bone formation or bone resorption responses. Our model explicitly considers how local mechanical signals propagate through the network before inducing such bone responses.

#### Evolution of bone boundaries

The boundary positions in the discrete model are updated over a time step Δ*t* based on the number of osteoblasts Ob_±_(*t*) and osteoclasts Oc_±_(*t*) at time *t* at the left boundary (indicated by −) and at the right boundary (indicated by +) according to:

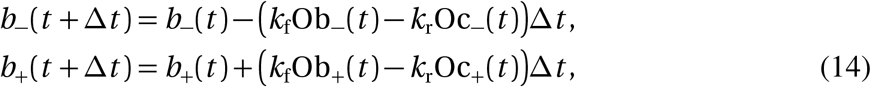

where *k*_f_ *>* 0 and *k*_r_ *>* 0 are the bone formation and bone resorption coefficients, respectively. These coefficients represent the length of bone produced or removed per cell per unit time, with units ofµm/day. In the continuum limit as Δ*t* approaches zero, Eqs (14) become

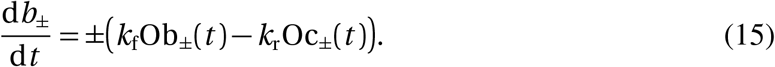

#### Osteoblast and osteoclast generation

We consider two different models for the generation of osteoblasts and osteoclasts at the bone boundaries:

1. *Instantaneous cell generation model*. In this variant, the number of osteoblasts and osteoclasts at each bone boundary relates directly to the difference in flux of signalling molecules at the bone surface compared to the reference value *J*_0_:

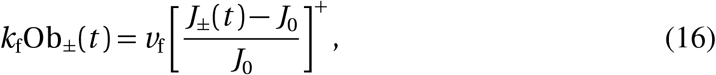

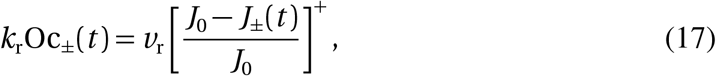

where *v*_f_ *>* 0 and *v*_r_ *>* 0 are bone formation and resorption speeds, respectively, with units ofµm/day, and [*j*] ^+^ = max {0, *j*} indicates taking the positive part. This model reflects common Wolff’s laws in which a bone apposition rate or bone resorption rate (inµm/day) is directly proportional to a mechanical stimulus, except that here this stimulus is a signal that propagates through the osteocyte network. We note here that it is not possible for the instantaneous cell generation model to have both osteoblasts and osteoclasts at the same boundary at a given time.
2. *Differential cell generation model*. In this model variant, it is the rate of change of the number of osteoblasts and osteoclasts at each bone boundary that relates to the difference in flux of signalling molecules at the bone surface compared to the reference value *J*_0_:

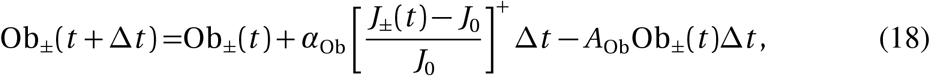

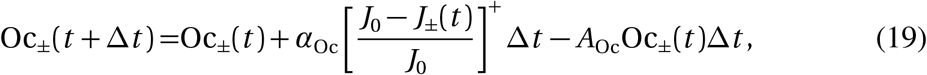

respectively. The rate parameters *α*_Ob_, *α*_Oc_ (in day^−1^) modulate the strength of osteoblast and osteoclast generation, and *A*_Ob_, *A*_Oc_ (in day^−1^) specify the elimination rates of osteoblasts and osteoclasts. In the continuum limit Δ*t* → 0, Eqs (18) and (19) become differential equations for the population of osteoblasts and osteoclasts:

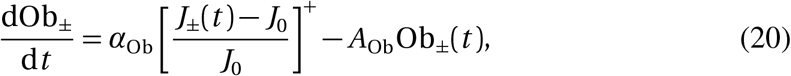

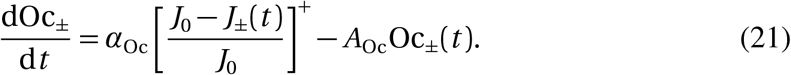

In contrast to the instantaneous cell generation model, the differential cell generation model allows both osteoblasts and osteoclasts to be present at the same boundary at a given time. While only one cell type is usually found in a given region of the bone surface (Martin, Burr & Sharkey 1998), the co-existence of osteoblasts and osteoclasts at the bone boundaries in our model represents an average over a small representative portion of the bone surface.

#### Osteocyte network evolution

During new bone formation, some bone forming cells become embedded into the new bone matrix and differentiate into osteocytes (Martin, Burr & Sharkey 1998). Osteocytes are usually found to be regularly spaced (Marotti 1992; Ardizzoni 2001). In our model, we assume that when new bone is formed, a new osteocyte is created during a time increment Δ*t* as soon as the bone surface extends a distance *Δx* from the osteocyte closest to the boundary (Fig. 1c). At the right boundary, this occurs when

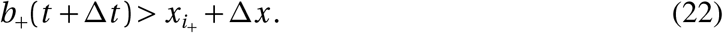

In this case, *i*_+_ is incremented by one, i.e., *i*_+_(*t* + Δ*t*) = *i*_+_(*t*) + 1. The newly formed osteocyte is placed at *i*_+_(*t* + Δ*t*)Δ*x* and initiated with zero signalling molecules. It will start producing signalling molecules based on the reaction term *R*_*σ*_(*x, t, N*) in Eq. (4) or *R*_*ψ*_(*x, t, N*) in Eq. (6).

An osteocyte is removed from the network during a time increment Δ*t* when bone resorption results in the bone surface moving past its position. At the right boundary, this occurs when

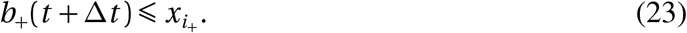

In this case, *i*_+_ is decremented by one, i.e., *i*_+_(*t* + Δ*t*) = *i*_+_(*t*) − 1. Signalling molecules that were associated with the resorbed osteocyte are assumed to be freed into the microenvironment and to contribute to *N*_+_(*t*) used in Eqs (2)–(3) to calculate the flux of signalling molecules at the bone surface sensed by osteoblasts and osteoclasts (see Figure 1d). Similar considerations hold for the formation and removal of osteocytes at the left boundary.

#### Numerical simulations

To simulate our discrete model numerically, molecular occupancies and the fluxes of signalling molecules emanating from the bone surfaces are first evolved using Eqs (1)–(5). Populations of osteoblasts and osteoclasts are then calculated using either Eqs (16)–(17) (instantaneous cell generation model), or Eqs (18)–(19) (differential cell generation model). Finally, bone boundaries are evolved with Eqs (14) and the osteocyte network is updated based on Eqs (22)–(23). The discrete model is initialised with no osteoblasts nor osteoclasts, and molecular occupancies corresponding to the steady state of the continuum limit, see Section 2.3, Eq. (25). The model is then first evolved to steady state under a force *F*, before changes to the loading force are applied. For full implementation details, the reader is referred to the computer code available on Github (Mehrpooya, Challis & Buenzli 2025). All parameter values of the model used in the numerical simulations are summarised in Table 1. Their estimation is detailed in Section 2.3.

The continuum equations derived from the discrete model form a reaction–diffusion problem for the concentration of molecules on an evolving domain. Mehrpooya, Challis & Buenzli (2024) provide an extensive comparison between discrete and continuum models of the propagation of signalling molecules within networks. Here, we use the continuum model mostly for parameter estimation and analysis.

### 2.3 Model parameter estimation

We now describe how we calibrate our mathematical model and select some of its parameters for biologically sensible behaviours.

We first consider the speed of transduced signal propagation through the osteocyte network to estimate the jump paramaters 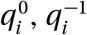, and 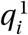. In our previous work (Mehrpooya, Challis & Buenzli 2024), we presented an application of signal propagation in bone with an inhomogeneous density of osteocytes and second-neighbour jumps. Here, we restrict to nearest-neighbour jumps and homogeneous osteocyte density, and therefore adjust the jump probabilities slightly compared to this previous work. We set 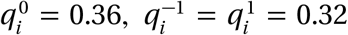 over a time step of Δ*t* = 10^−5^ day for all *i*. In the continuum limit in Eqs (11)–(12), this corresponds to *D* = 3.2 mm^2^/day ≈ 37µm^2^/s and *v* = 0. These values lead to reasonable rates of signal propagation through the osteocyte network, with a diffusive root mean square displacement of 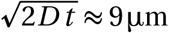 over one second, comparable to calcium signal travelling speeds of 2µm/s (Turner & Forwood 1995) and fluid flow velocities of 20 − 60µm/s (Verbruggen, Vaughan & McNamara 2014; van Tol et al. 2020b).

The reaction rate parameters *λ*_1_ and *λ*_2_ influence the number of signalling molecules produced by osteocytes and emanating from the bone surface to induce bone formation or bone resorption. Some changes are required as well for values assumed for *λ*_1_ and *λ*_2_ compared to our previous work (Mehrpooya, Challis & Buenzli 2024), because we now include mechanics-dependent production rates to model osteocyte mechanotransduction in Eqs (4), (6). To ensure that bone adaptation to loading forces is consistent with experimental observations, we proceed in two steps. In a first step, we consider homeostatic steady states in which bone length *L* is adapted to a constant applied loading force *F*. These steady states provide a useful relationship *L*(*F*) that enables us to estimate the reaction rate parameters *λ*_1_, *λ*_2_, and the threshold flux value *J*_0_. In a second step, we consider the dynamics of bone adaptation leading to these homeostatic steady states, which enables us to estimate parameters influencing osteoblast and osteoclast populations.

#### Homeostatic steady states: bone length under constant loading force

To understand how *λ*_1_, *λ*_2_ and *J*_0_ influence homeostatic steady states in which bone length is stationary under a constant loading force, we reason on the continuum model and determine the relationship *L*(*F*) between an applied constant force *F* and the corresponding homeostatic bone length *L*. This kind of relationship is typical in Wolff’s laws, in which the bone steady state is in a one-to-one relationship with the mechanical forces exerted on bone (Turner 1999; Lerebours & Buenzli 2016).

In steady state, *∂n/∂ t* = 0 in Eq. (10) and d*b*_±_/d*t* = 0 in Eq. (15). With the stress-dependent reaction rate from Eq. (4), Eq. (10) becomes

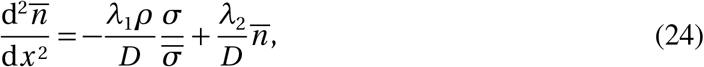

where 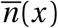 is the steady-state concentration profile of signalling molecules within bone, and *σ* is constant and uniform due to the constant loading force, see Eq. (5). Equation (24) is an inhomogeneous second-order differential equation with constant coefficients. Since there is no bone boundary motion, we can always choose *b*_±_ = ±*L/*2 for symmetry, where *L* is the fixed bone length. The solution to Eq. (24) with Dirichlet boundary conditions (Eq. (13)) at *x* = ±*L/*2 is then given by

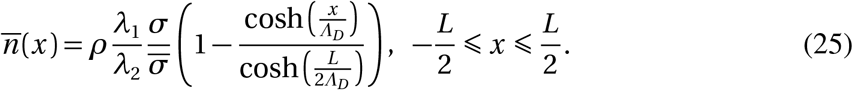

In Equation (25),

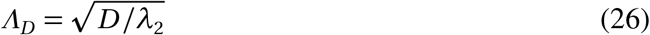

is the diffusion length of the signalling molecules, which represents the average distance that signalling molecules travel by diffusion during their lifetime 1*/λ*_2_. The parameter *Λ*_*D*_ controls the relative strength of diffusive transport over reaction. When *Λ*_*D*_ is large compared to the bone length *L*, transport dominates reaction and the continuum limit represents the discrete model well (Simpson et al. 2010; Li et al. 2022; VandenHeuvel, Buenzli & Simpson 2024; Mehrpooya, Challis & Buenzli 2024). The steady-state boundary fluxes 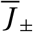 are given by

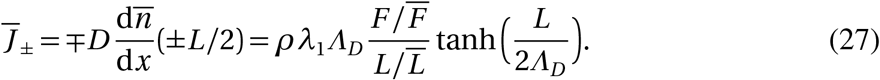

From Eq. (15), osteoblast and osteoclast populations in steady state, denoted by 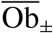 and 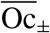, respectively, are such that 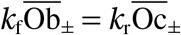. Therefore, with the instantaneous cell generation model (Eqs (16) and (17)), the steady-state boundary fluxes 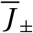 must satisfy

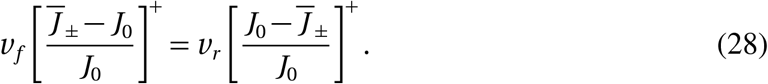

With the differential cell generation model (Eqs (20) and (21)), the steady-state boundary fluxes 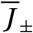 must satisfy

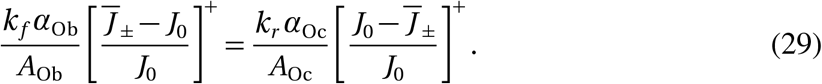

The only way to satisfy these equations is to have 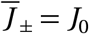, which implies also that 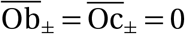 in both models. Therefore, in steady state, the flux of signalling molecules emanating from the bone surface must equal the threshold value *J*_0_, as expected, so that with Eq. (27),

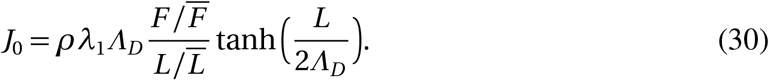

We can recast Eq. (30) to obtain an explicit relationship *F*(*L*) between the applied loading force *F* and the homeostatic bone length *L*, which depends on the choice of several parameters:

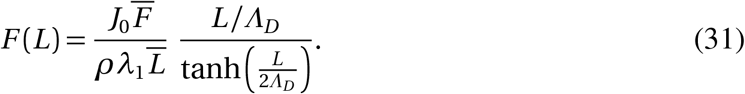

Inverting this equation determines an effective steady-state Wolff’s law *L*(*F*). Figure 2 shows the curve *F*(*L*) plotted for different values of *Λ*_*D*_ when all curves go through the point 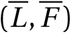, which occurs when the parameters satisfy

**Figure 2.**
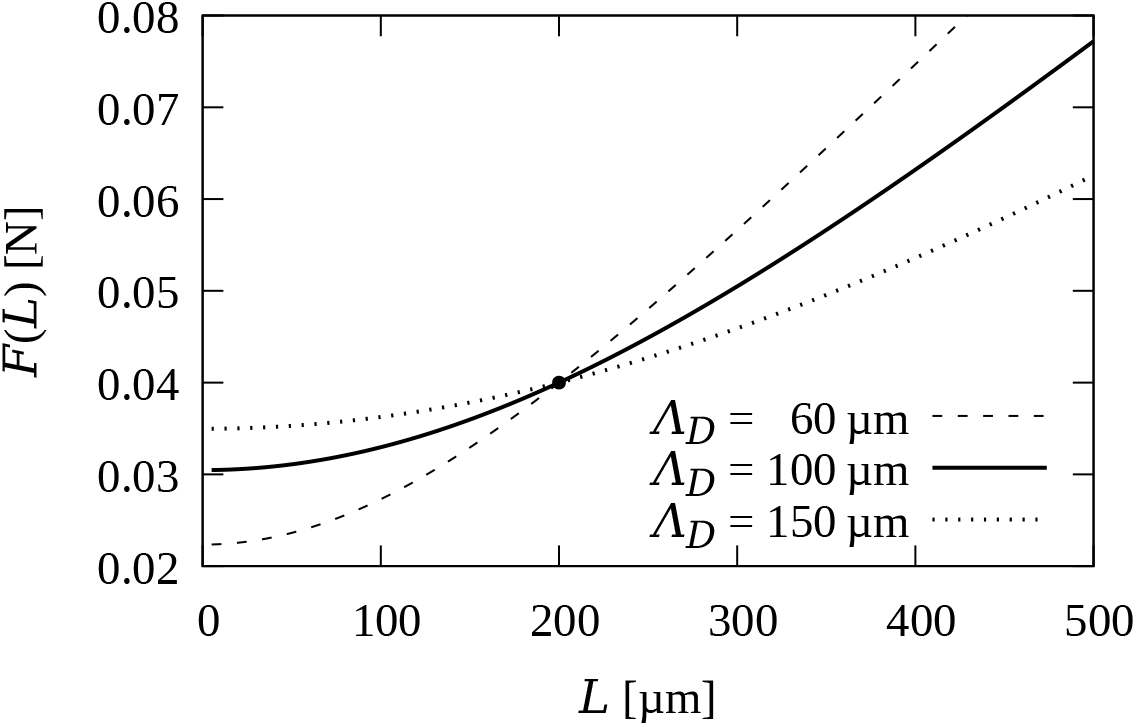
Relationship between loading force *F* and homeostatic bone length *L* with 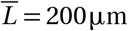 and 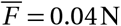 in Eq. (33) for *Λ*_*D*_ = 60µm, 100µm, and 150µm

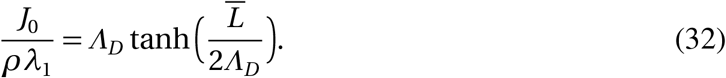

In this case, Eq. (31) can be recast as

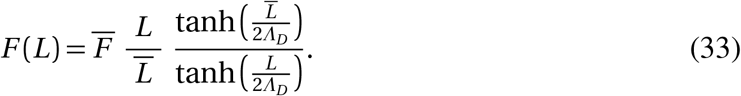

In this expression for *F*(*L*), the diffusion length *Λ*_*D*_ affects both the steepness of *F* ≫ (*L*) for *L ≫ Λ*_*D*_, and the width of the region 0 ⩽ *L* ≲ *Λ*_*D*_ in which *F*(*L*) does not vary significantly (Figure 2). Given our choice of reference mouse tibia cortical wall width 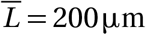, we set *Λ*_*D*_ = 100µm. This value of *Λ*_*D*_ is large enough for diffusion-dominated signal propagation so the continuum limit matches the discrete model, yet small enough so that small changes in *F* do not cause excessive changes in *L*.

Having set *D* = 3.2 × 10^6^µm^2^/day and *Λ*_*D*_ = 100µm, the degradation rate of signalling molecules is determined from Eq. (26) to be *λ*_2_ = 320/day. From Eq. (32) with *ρ* = 1/Δ*x* = 0.1µm^−1^, we must also have *J*_0_*/λ*_1_ = 10 tanh(1) ≈ 7.62, and so we finally set *λ*_1_ = 128/s = 11, 059, 200/day, and *J*_0_ = 84226220/day. The value of *λ*_1_ scales the number of signalling molecules at osteocytes in steady state. Here it is chosen to be slightly larger than values used in our previous work (Mehrpooya, Challis & Buenzli 2024).

As illustrated above, the continuum limit of the discrete model is particularly useful to reason qualitatively and quantitatively on the behaviour of the discrete model. We have used these behaviours to determine key parameter values associated with signal propagation and steady-state configurations so they are consistent with expected bone adaptation properties. To ensure that these properties carry over to the discrete model, we assess in Figure 3 the discrepancy between the discrete model and its continuum limit in steady state using the parameters estimated so far. We compare the steady state of the discrete model for a fixed bone length of 200µm and a uniform loading force 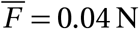 with the continuum steady state 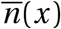 from Eq. (25). Figure 3 shows that there is a good match between the steady state molecule concentration profiles of the discrete and continuum models (grey histogram bars and dashed red curves, respectively), and a good match between the continuum and discrete boundary fluxes (black and red slopes). The flux discrepancy between the discrete and continuum models is about 7.5%.

**Figure 3.**
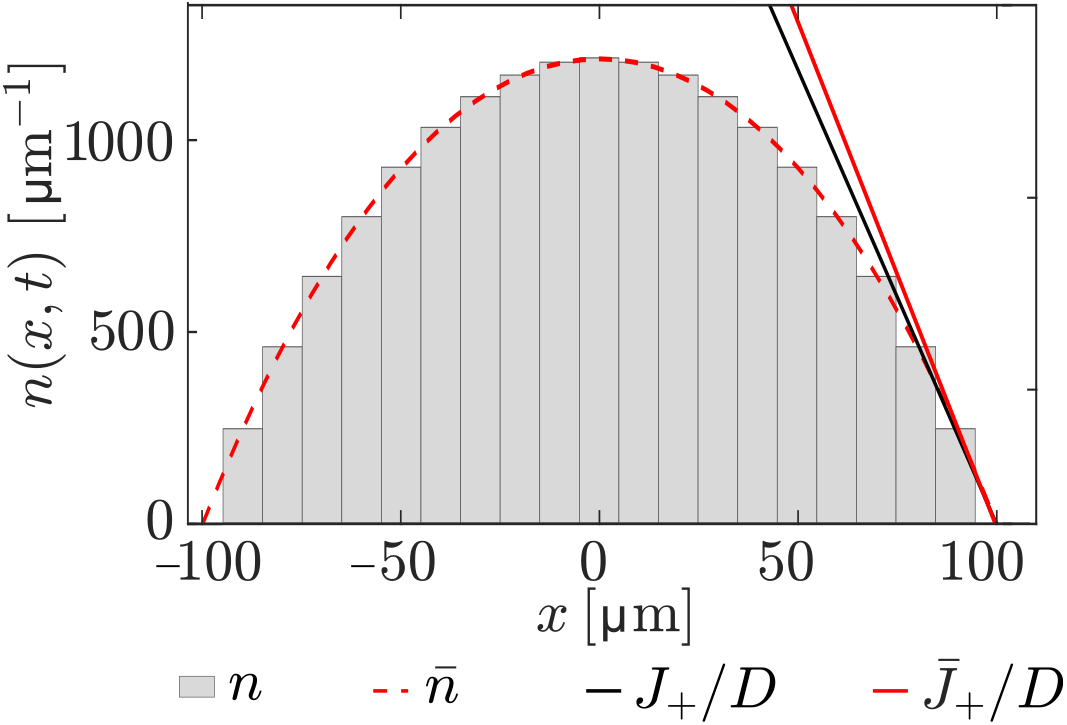
Comparison of the steady state calculated from the discrete and continuum models with fixed boundaries and uniform loading force. Molecule concentrations corresponding to the discrete (grey) and exact (red dashed) solutions at steady state for a bone length of 200µm. The solid black line represents the slope *∂n/∂ x* = −*J*_+_*/D* of the discrete concentration at the right boundary, calculated as *n*_*i*+_/Δ*x*. The solid red line represents the slope of the continuum concentration at the right boundary, calculated as *∂n/∂ x*(*L/*2) from Eq. (25)

#### Osteoblast and osteoclast parameters

The remaining parameters of the model, *k*_f_, *k*_r_, and either *v*_f_ and *v*_r_ (instantaneous cell generation model) or *α*_Ob_, *α*_Oc_, *A*_Ob_, *A*_Oc_ (differential cell generation model) are associated with the population of osteoblasts and osteoclasts. These parameters are estimated in Appendix A based on the dynamics of bone adaptation to steady state under a 20% increase in the force. In brief, the instantaneous cell generation model with moving boundaries leads to biologically unrealistic instabilities in the form of oscillations of increasing amplitude in the flux of signalling molecules and in boundary positions (Section A.1). These oscillations are not overcome by tuning *v*_f_ and *v*_r_. We therefore do not consider the instantaneous cell generation model further, except in the analysis of bone behaviour under a force gradient in Section 3.4. The instabilities of the instantaneous cell generation model are resolved by the differential cell generation model, which takes into account the fact that the generation of osteoblasts and osteoclasts in response to mechanical signals takes some time. While oscillations in boundary positions may still occur for some parameter combinations in the differential cell generation model, these oscillations are stable, and a parameter regime can be devised in which these oscillations are overdamped (Section A.2).

## 3 Results and discussion

In this section we first determine steady states of the discrete model with moving boundaries reached by initialising it with steady states of the continuum model for a variety of bone lengths *L* and loading forces *F*. We then present a range of numerical results that demonstrate key features of our discrete model for bone mechanical adaptation under temporal and spatial variations of the loading force.

Figure 4 highlights the difference in steady-state lengths obtained for constant loading forces in the continuum model (red curve, given by Eq. (31)) and in the discrete model (blue crosses). The discrete simulations are initialised from the continuum steady states by setting signalling molecule occupancies as 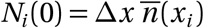, where 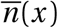 is the steady-state concentration of the continuum model in Eq. (25) for various initial lengths *L* (0). The time evolution of the discrete model is illustrated in this figure by trajectories *L*(*t*), *F* (blue lines) ending at the discrete model steady state (blue crosses). Overall there is a good match between the steady-state lengths obtained with the discrete and continuum models. However, at all values of the applied loading force, the equilibrium states of the discrete model have a slightly shorter bone length compared to the equilibrium states predicted by the continuous model. The match improves at lower values of *L*, as expected based on the continuum limit being valid for *Λ*_*D*_ ≫ *L* (VandenHeuvel, Buenzli & Simpson 2024; Mehrpooya, Challis & Buenzli 2024). In our subsequent numerical simulations, we initialise the discrete model with the homeostatic steady state 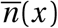 obtained from the continuum model with 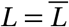, but always wait until the discrete model reaches a steady state before varying the loading force.

**Figure 4.**
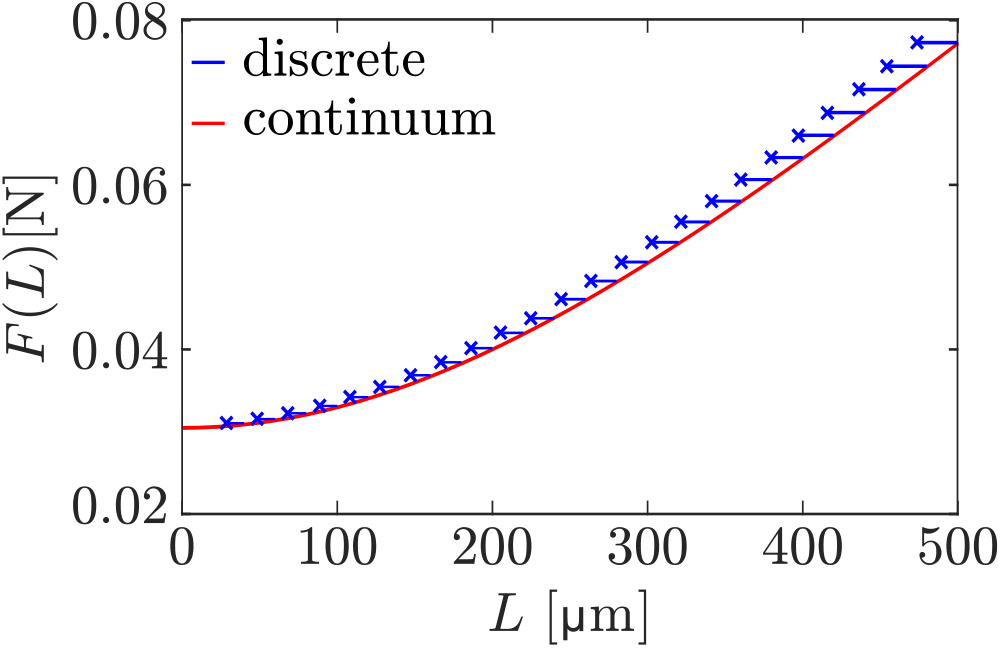
Discrete model trajectories of the bone length (blue lines) alongside the continuum steady-state bone length (red curve) for different uniform loading forces and initial network lengths

### 3.1 Bone response to overloading followed by unloading

We now examine the response of the discrete model under a cycle of overloading and unloading. We assume that the steady-state force 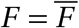 is first uniformly increased by 20% at *t* = 40 days, from *F* = 0.04 N to *F* = 0.048 N, and then restored to its initial value *F* = 0.04 N at *t* = 200 days.

Snapshots of the concentration profiles *n*(*x*_*i*_, *t*) at each osteocyte location at various times are shown in Figure 5 (grey bars) and compared with the steady-state concentration profiles 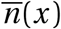 from Eq. (25), where *L* is taken as *L* = (*i*_+_ + 1)Δ*x* − (*i*_−_ − 1)Δ*x* and *σ* is taken as *σ*(*t*) = *F*(*t*)/[(*b*_+_(*t*) − *b*_−_(*t*)) *W*](red dashed line). There is an excellent match between the discrete model concentrations and the continuum profiles, despite the continuum profiles assuming a steady state, indicating that the dynamics of molecular signalling is fast compared to the dynamics of bone changes. The goodness of the match relies on assuming that 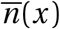 has absorbing boundary conditions at (*i*_−_−1)Δ*x* and (*i*_+_ + 1)Δ*x* (red dotted vertical lines) instead of *b*_−_ and *b*_+_ (blue vertical lines), respectively.

**Figure 5.**
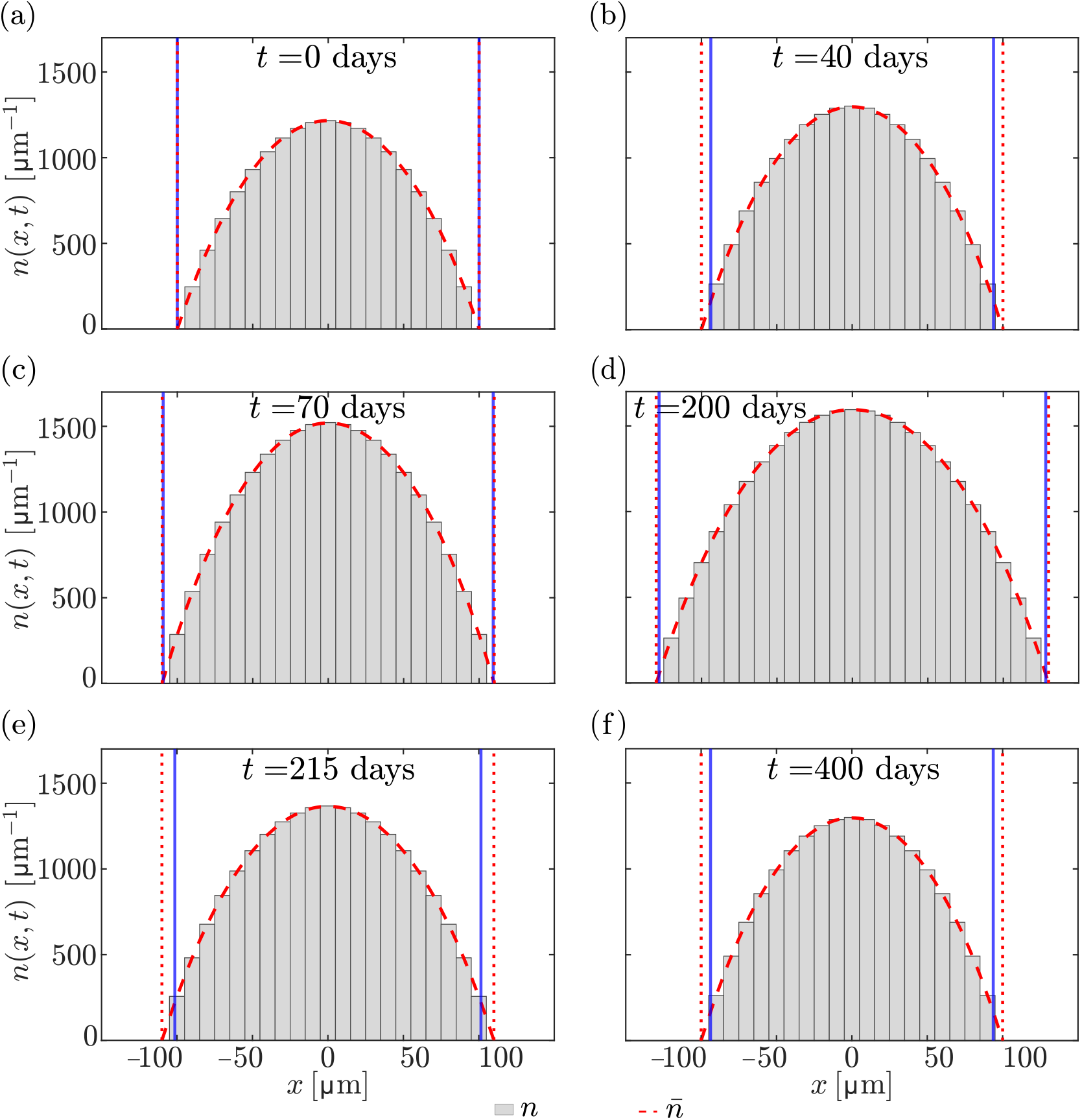
Snapshots of concentration profiles *n*(*x*_*i*_, *t*) (grey bars) and steady-state concentration profiles 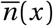 from Eq. (25) (dashed red line) with absorbing boundary conditions at (*i*_−_− 1)Δ*x* and (*i*_+_ + 1)Δ*x* (red dotted vertical lines), shown at various times. Vertical blue lines indicate the bone boundaries *b*_−_ (*t*) and *b*_+_(*t*). (a) Initial condition *t* = 0; (b) Steady state of the discrete model, *t* = 40 days; (c) Intermediate state during the formation phase, *t* = 70 days; (d) New steady state reached after the force increase, *t* = 200 days; (e) Intermediate state during the resorption phase, *t* = 215 days; (f) Final steady state after return to the initial loading force, matching the state at *t* = 40 days

Figure 6a illustrates the time evolution of the relationship between bone length and mechanical force by plotting the trajectory *L*(*t*), *F*(*t*) (blue). This trajectory begins at 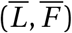 on the *F*(*L*) curve given by Eq. (31) (red). It first evolves to 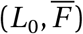 (blue cross), where *L*_0_ denotes the steady-state bone length in the discrete model for the force 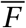. After the force is increased at *t* = 40 days, bone length increases and the trajectory tends toward the *F*(*L*) curve with a higher value of *F*. After restoring the force to its initial value 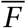 at *t* = 200 days, bone length returns to its steady-state bone length *L*_0_.

**Figure 6.**
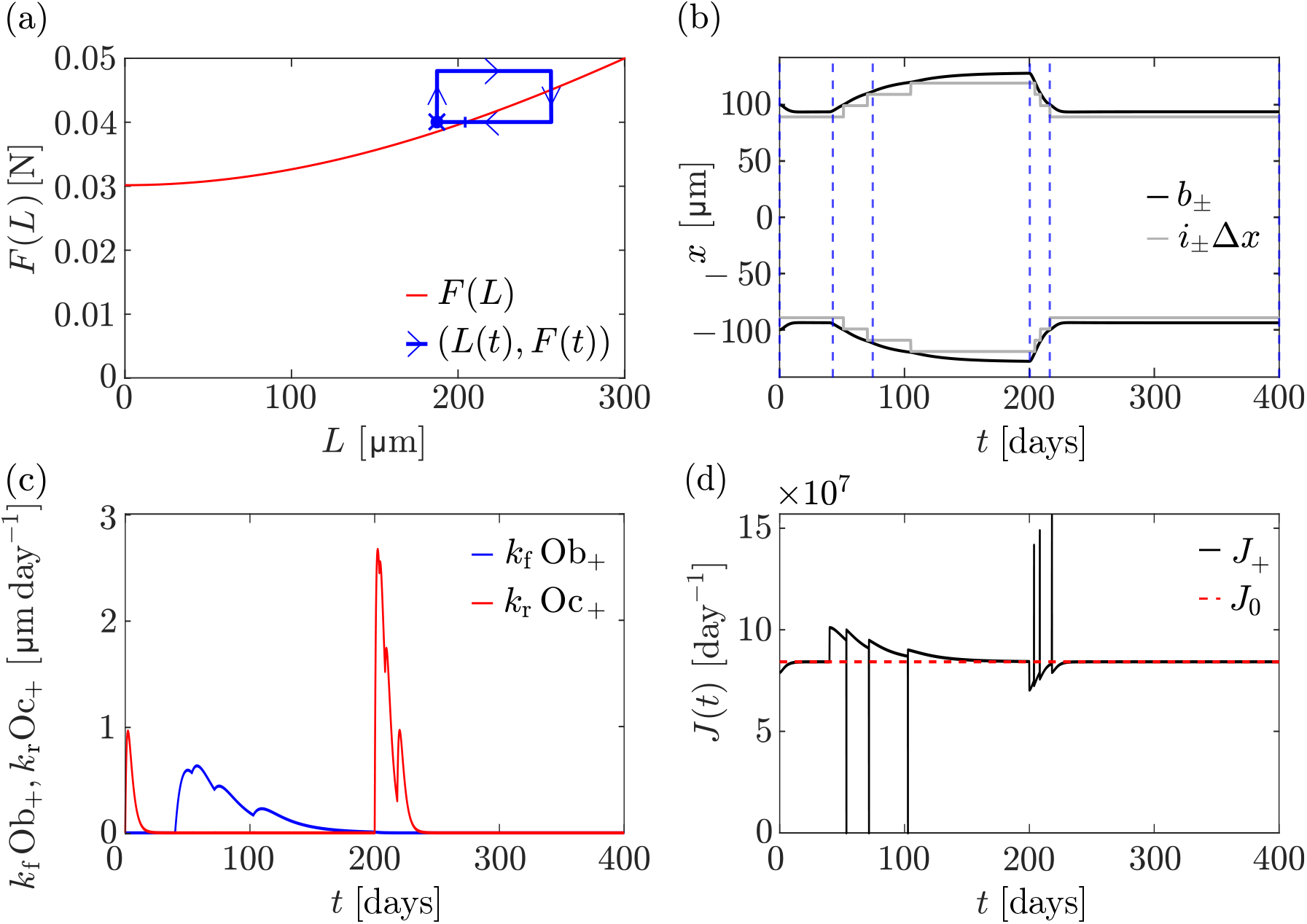
Overloading followed by unloading. At *t* = 40 days, a 20% force increase is applied, from *F* = 0.04 N to *F* = 0.048 N, followed by a return to *F* = 0.04 N at *t* = 200 days. (a) The trajectory *L*(*t*), *F*(*t*) from the discrete model simulation (blue line) is shown alongside the relationship *F*(*L*) in Eq. (33) between bone length and mechanical force in the homeostatic steady state of the continuum model (red line). The trajectory starts on the *F*(*L*) curve (blue vertical bar), converges to a discrete steady state 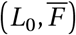 at *t* = 40 days (blue cross), consistent with Figure 4. The final point of the trajectory at *t* = 400 days (blue dot) coincides with the steady state 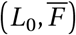 at *t* = 40 days. (b) Evolution of *i*_±_ Δ*x* (grey lines), along with the left and right boundary positions (black lines) over time. Blue dashed vertical lines denote time points at which snapshots of the molecule concentrations are shown in Figure 5. (c) Speed of osteoblast-driven (blue line) and osteoclast-driven (red line) boundary movement at the right boundary over time. (d) Evolution of the flux at the right boundary for the discrete model (black line), with the reference value *J*_0_ shown (red dashed line)

The force increase at *t* = 40 days first leads to an increase in the production of signalling molecules within the osteocyte network, which increases their concentration (Figure 5b–d). This surplus of signalling molecules propagates to the bone surface and induces an increase in boundary fluxes *J*_±_, which in turn stimulates osteoblast generation (Figure 6c) and bone growth (Figure 6b). The formation of osteocytes in this overloading phase is marked by steps in the evolutions of *i*_±_(*t*)Δ*x* in Figure 6b. When a new osteocyte is formed at the growing bone boundary, it has no signalling molecules. This induces a sudden, but short-lived decrease in the boundary fluxes (Figure 6d), until the new osteocyte collects signalling molecules from its neighbour and from its own production. Bone length continues to increase until the boundary fluxes *J*_±_ converge toward the reference value *J*_0_.

The evolution of the population of osteoclasts and osteoblasts and of the flux of signalling molecules emanating from the bone surfaces during the overloading and unloading cycle are shown in Figures 6c and 6d, respectively. The population of osteoclasts at early times *t <* 40 days is generated due to the flux *J*_±_ being initially slightly lower than the reference value *J*_0_. This difference is due to *J*_0_ being estimated from the steady state of the continuum limit while *J*_±_ are calculated from the discrete model (see Section 2.3 and Figure 3). This initial bone resorption explains why the steady-state length of the discrete model is slightly less than that predicted from the continuum model by the *F*(*L*) curve.

Interestingly, during this formation phase, the pointwise generation of osteocytes and its effect on boundary fluxes disrupts the population of osteoblasts, which exhibits several small peaks (Figure 6c). It is possible that these kinds of discrete behaviours could be related to the formation of regular spatial patterns seen in lamellar bone, in which bone is produced in distinct layers called lamellae, with possibly layered osteocyte generation (Martin, Burr & Sharkey 1998; Marotti 1992; Ardizzoni 2001).

When the force is restored at *t* = 200 days, osteocytes within the network decrease their production of signalling molecules. This depletion of signalling molecules propagates to the bone surface where it induces a reduction in boundary fluxes *J*_±_, which in turn stimulates osteoclast generation (Figure 6c) and leads to bone loss (Figure 6b). Osteocyte resorption is marked by steps in the evolutions of *i*_±_(*t*)Δ*x*. When an osteocyte is resorbed, its signalling molecules are freed into the micro-environment. This generates a strong, but short-lived spike in the boundary fluxes (Figure 6d). The steady state is reached when the boundary fluxes converge to the reference value *J*_0_.

### 3.2 Bone response to unloading followed by reloading

We now examine the model’s behaviour when unloading occurs before reloading. We assume that the steady-state force 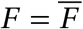 is first uniformly reduced by 20% at *t* = 40 days, i.e., from *F* = 0.04 N to *F* = 0.032 N, and then restored to its initial value *F* = 0.04 N at *t* = 200 days.

The trajectory *L*(*t*), *F*(*t*) in Figure 7a (blue) shows a similar pattern as in Figure 6a, where there is first a period of convergence to the steady state bone length *L*_0_ of the discrete model. When the force is reduced at *t* = 40 days, bone length decreases and the trajectory tends toward the *F*(*L*) curve at the reduced value of *F*. After restoring the force to its initial value 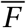 at *t* = 200 days, bone length increases and converges to a value *L*_final_. However, here, the final bone length *L*_final_ does not converge back to the initial steady-state length *L*_0_ of the discrete model. The trajectory is not closed, meaning that bone length is not fully recovered after the force is restored to its initial value. This is also visible from the evolution of boundary positions *b*_±_(*t*) in Figure 7b.

**Figure 7.**
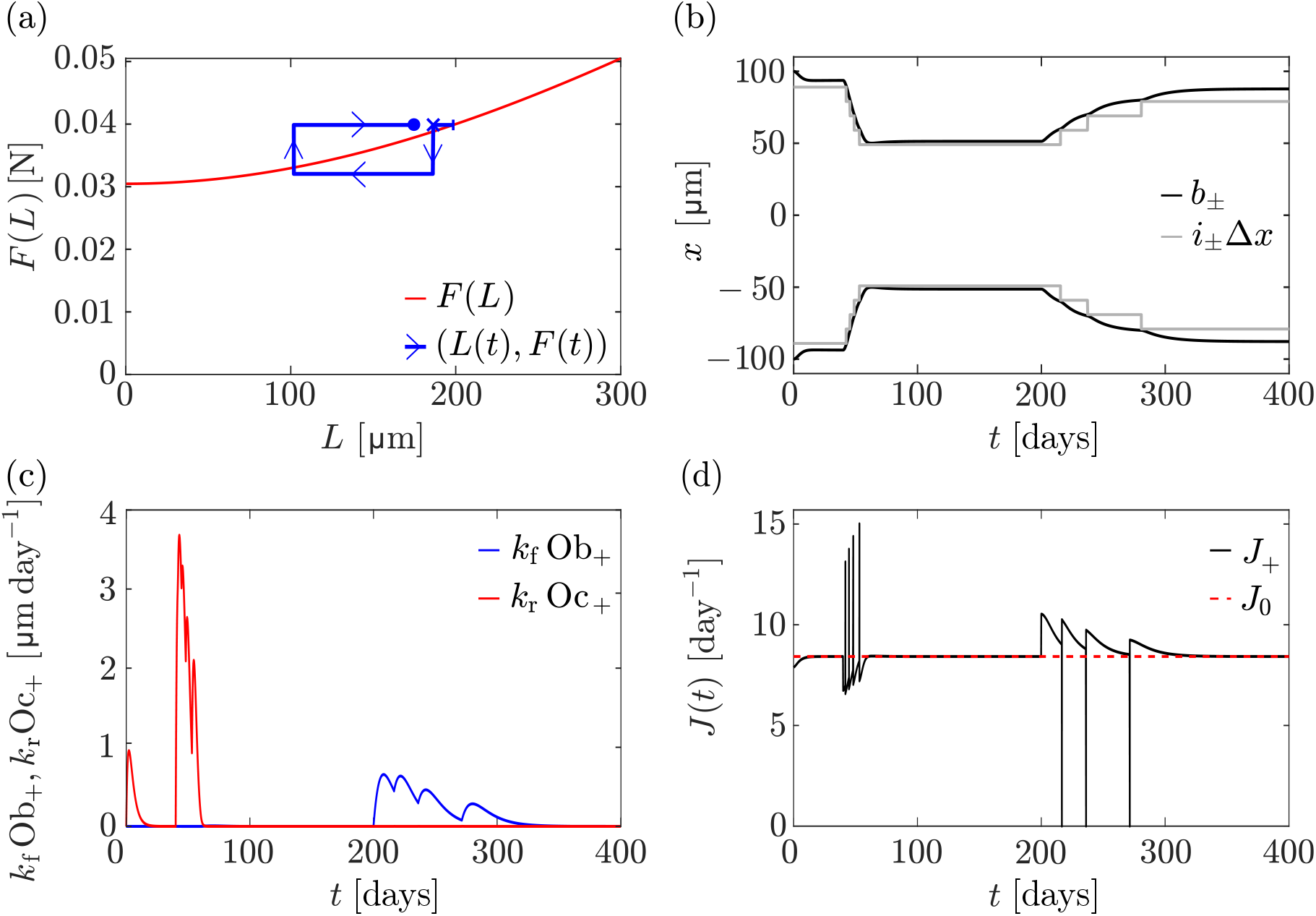
Unloading followed by reloading. At *t* = 40 days, a 20 % force reduction is applied, from *F* = 0.04 N to *F* = 0.032 N, followed by a return to *F* = 0.04 N at *t* = 200 days. (a) The trajectory *L*(*t*), *F*(*t*) from the discrete model simulation (blue line) is shown alongside the relationship *F*(*L*) in Eq. (33) between bone length and mechanical force in the homeostatic steady state of the continuum model (red line). The trajectory starts on the *F*(*L*) curve (blue vertical bar), converges to a discrete steady state 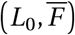 at *t* = 40 days (blue cross), consistent with Figure 4. The final point of the trajectory at *t* = 400 days (blue dot) is a new steady state 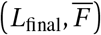 that does not coincides with the steady state 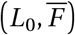 at *t* = 40 days. (b) Evolution of *i*_±_Δ*x* (grey lines), along with the left and right boundary positions (black lines) over time. (c) Speed of osteoblast-driven (blue line) and osteoclast-driven (red line) boundary movement at the right boundary over time. (d) Evolution of the flux at the right boundary for the discrete model (black line), with the reference value *J*_0_ shown (red dashed line)

Figure 7b shows that while four osteocytes are resorbed at either ends during the unloading phase, only three are re-generated during the reloading phase. This behaviour shows that history dependence of the bone state may arise due to the discrete nature of osteocytes accounted for in our discrete model. Such history dependence is not possible in the continuum model, where steady-state lengths under a constant uniform force are always unique, and characterised by the curve *F*(*L*).

The evolution of the population of osteoclasts and osteoblasts and of the boundary fluxes of signalling molecules exhibit similar qualitative patterns as in Figure 6. The temporary increases in osteoblasts after each osteocyte generation are more pronounced than in Figure 6. This may be due to bone possessing a lower number of osteocytes, and therefore being more strongly affected by an osteocyte formation. The fact that bone length is not fully restored after the force is returned to its initial value may also be an effect of small osteocyte numbers.

It is well-established experimentally that loading history plays an important role for the mechanical adaptation of bone (Schriefer et al. 2005; Turner 1999; Wang, Fu & Yang 2025). Incomplete recovery of bone after unloading and reloading is observed in astronauts after they return from a long-term mission in space (Vico et al. 2000; Dudley-Javoroski et al. 2012; Edwards and Schnitzer 2015), as well as in controlled animal experiments (Cunningham et al. 2018, 2023). In experiments of bone adaptation in mice, a more diminished recovery after reloading was observed in older mice and in osteocyte-deficient mice (Cunningham et al. 2023). These observations are consistent with our model results, where the availability of fewer osteocytes during reloading leads to incomplete bone recovery.

### 3.3 Disuse

The force–length relationship *F*(*L*) in Eq. (31) shows that the steady-state bone length goes to zero for a loading force *F* (0) ≈ 0.0304 N (Figure 2). This suggests that *F* (0) is a threshold force below which all bone is resorbed. In Figure 8, we simulate a disuse situation in which the loading force is reduced below the threshold to *F* = 0.02 N at *t* = 20 days. Contrary to the 20% unloading situation of Figure 7, here the bone is fully resorbed. The initial decrease in boundary fluxes at the onset of disuse generates osteoclasts that decrease bone length. Mechanical stresses in Eq. (5) increase with time due to the loss of bone, which stimulates signalling molecule production to compensate their initial depletion. However, these signalling molecules are produced by a decreasing number of osteocytes and the boundary fluxes of signalling molecules remain below the reference value *J*_0_ at all times until all the bone is resorbed.

**Figure 8.**
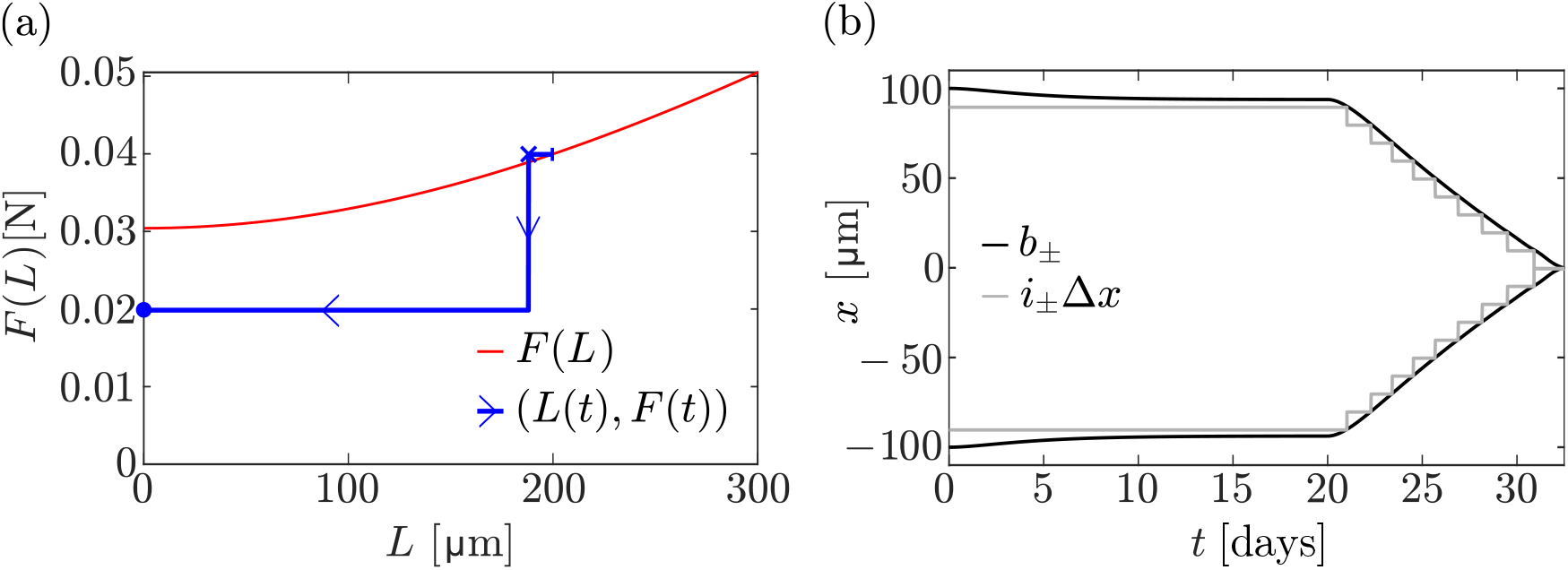
Disuse with stress stimulation of signalling molecule production. At *t* = 20 days, the force is reduced to *F* = 0.02 N, which is below the predicted minimum mechanical force threshold. (a) The trajectory *L*(*t*), *F*(*t*) of the discrete model simulation (blue line) is shown alongside the relationship *F*(*L*) in Eq. (33) (red line). The following points of the trajectory are marked: initial condition (blue vertical bar), steady state at *t* = 40 days (blue cross), final state (blue dot); (b) Evolution of *i*_±_Δ*x* (grey lines), along with the left and right boundary positions (black lines) over time

The destructive competition between the increase in signalling molecule production and the loss of osteocytes in this disuse situation can be overcome if a different mechanotransduction model is chosen, so that molecule production increases faster as bone is lost. This occurs, for example, if we assume that osteocytes produce signalling molecules in response to strain energy density *Ψ*(*x, t*) in Eqs (6)–(7). In this case, the relationship in Eq. (30) between the reference flux *J*_0_, the loading force *F*, and the bone length *L* in homeostatic steady states becomes

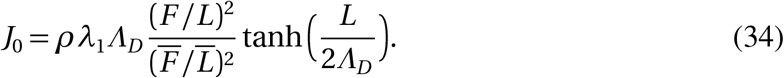

The relationship between loading force and homeostatic bone length found from Eq. (34) changes to

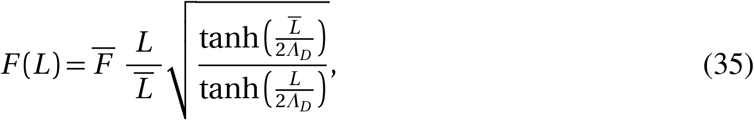

where parameters are chosen as in Eq. (32) so that 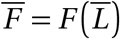. For small bone lengths, Eq. (35) shows that 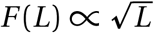, or, equivalently, *L*(*F*) ∝ *F* ^2^. This relationship is more protective of bone loss for small forces, and total bone loss is only obtained when *F* = 0. This is illustrated in Figure 9, in which the same disuse situation as in Figure 8 is simulated, but with a strain energy density stimulation of signalling molecule production by osteocytes (Eqs (6)–(7)). In this case, the production of signalling molecules increases quadratically with 1*/L* as bone length reduces. This increase in production rate is strong enough to compensate for the loss of osteocytes, and bone length reaches a nonzero steady state (Figure 9b).

**Figure 9.**
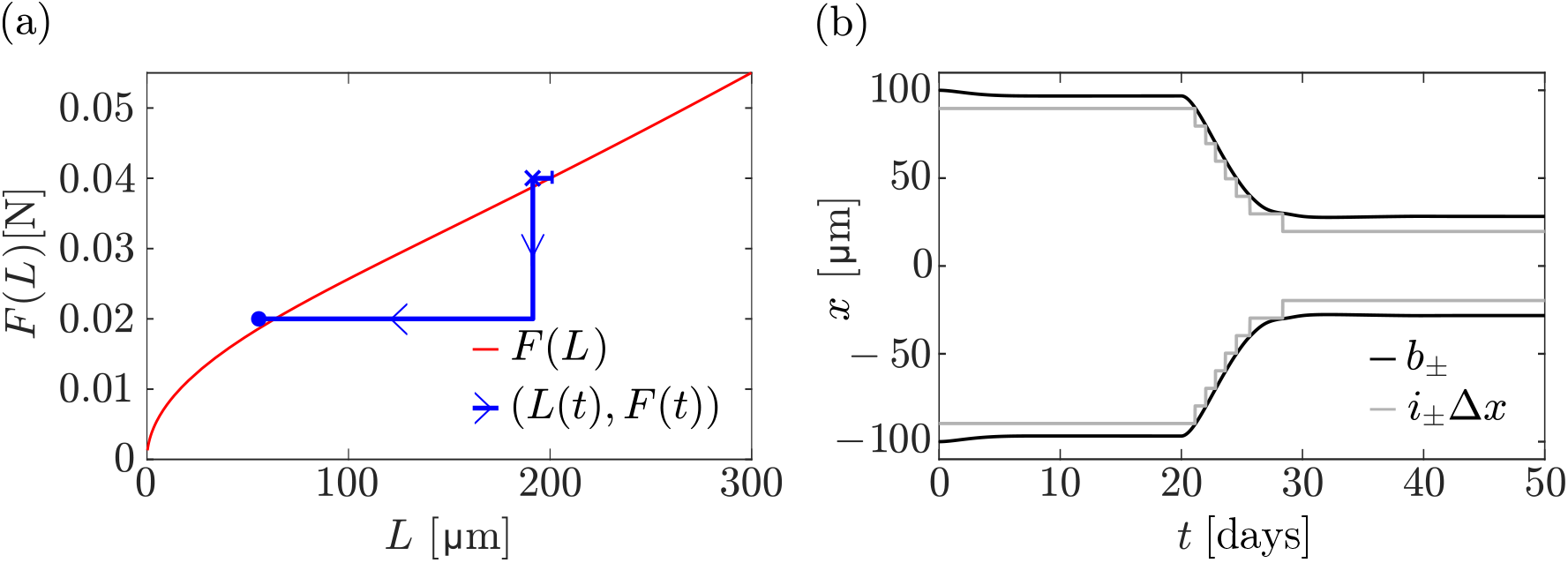
Disuse with strain-energy-density stimulation of signalling molecule production. At *t* = 20 days, the force is reduced to *F* = 0.02 N. (a) The trajectory *L*(*t*), *F*(*t*) of the discrete model simulation (blue line) is shown alongside the relationship *F*(*L*) in Eq. (35) (red line). The following points of the trajectory are marked: initial condition (blue vertical bar), steady state at *t* = 20 days (blue cross), final steady state at *t* = 50 days (blue dot); (b) Evolution of *i*_±_Δ*x* (grey lines), along with the left and right boundary positions (black lines) over time

The difference in behaviour of our model when osteocyte mechanotransduction is based on mechanical stress or when it is based on strain energy density is significant in a disuse situation. Wolff’s laws and similar mechanostat models predict that disuse leads to full resorption of bone, but this is only predicted for a zero loading force (Turner 1999; Skerry 2008). Our model shows that accounting for signal propagation through the osteocyte network enables new behaviours.

Disuse due to prolonged bedrest, spinal cord injury, and spaceflight missions shows that bone loss slows down and does not lead to complete loss. However, protective mechanisms such as osteocyte accommodation are thought to be at play in these situations (Turner 1999; Lerebours & Buenzli 2016). Our model does not account for such mechanisms. The existence of a threshold loading force below which all bone is resorbed in our model may not represent bone behaviour at a macroscopic scale. However, it could represent a tipping mechanism at a small scale during the structural optimisation of trabecular bone, by which trabecular struts are entirely resorbed as soon as the load they carry falls below a threshold (Kinney & Ladd 1998; Smotrova & Silberschmidt 2024).

### 3.4 Mechanical adaptation to a force gradient

We now investigate how our model predicts the mechanical adaptation of bone under a inhomogenous loading force

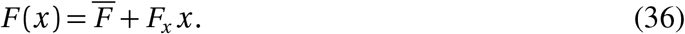

This force field can be thought of as that generated under compression (baseline 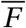) and bending (gradient *F*_*x*_) over one side of the cortical wall (Figure 1a). For this simulation, we use the stress-induced mechanostransduction model (Eqs (4)–(5)) and choose a gradient value *F*_*x*_ = 5 × 10^−5^ N*/µ*m, so that *F* (−100µm) = 0.035 N, which is slightly over the homogeneous loading force threshold 0.0304 N below which bone is fully resorbed (see Section 3.3). We initialise molecule concentrations as before, based on the homeostatic steady-state concentration 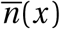 of the continuum model when 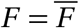, given by Eq. (25).

Figure 10 shows that under this force gradient, bone is first resorbed quickly at the left boundary because the force value there is less than 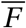. At the same time, bone is formed at the right boundary because the force value there is greater than 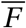. Therefore, the bone drifts in space toward greater values of *x* (Figure 10c). The speed at the right boundary is also greater than the speed at the left boundary, which results in an approximately linearly increasing bone length at large times (Figure 10d). The sequential removal of osteocytes at the left boundary and formation of new osteocytes at the right boundary leads to regular spikes in the boundary fluxes (Figure 10b), which generate sizeable fluctuations in the osteoclast and osteoblast populations (Figure 10e,f).

**Figure 10.**
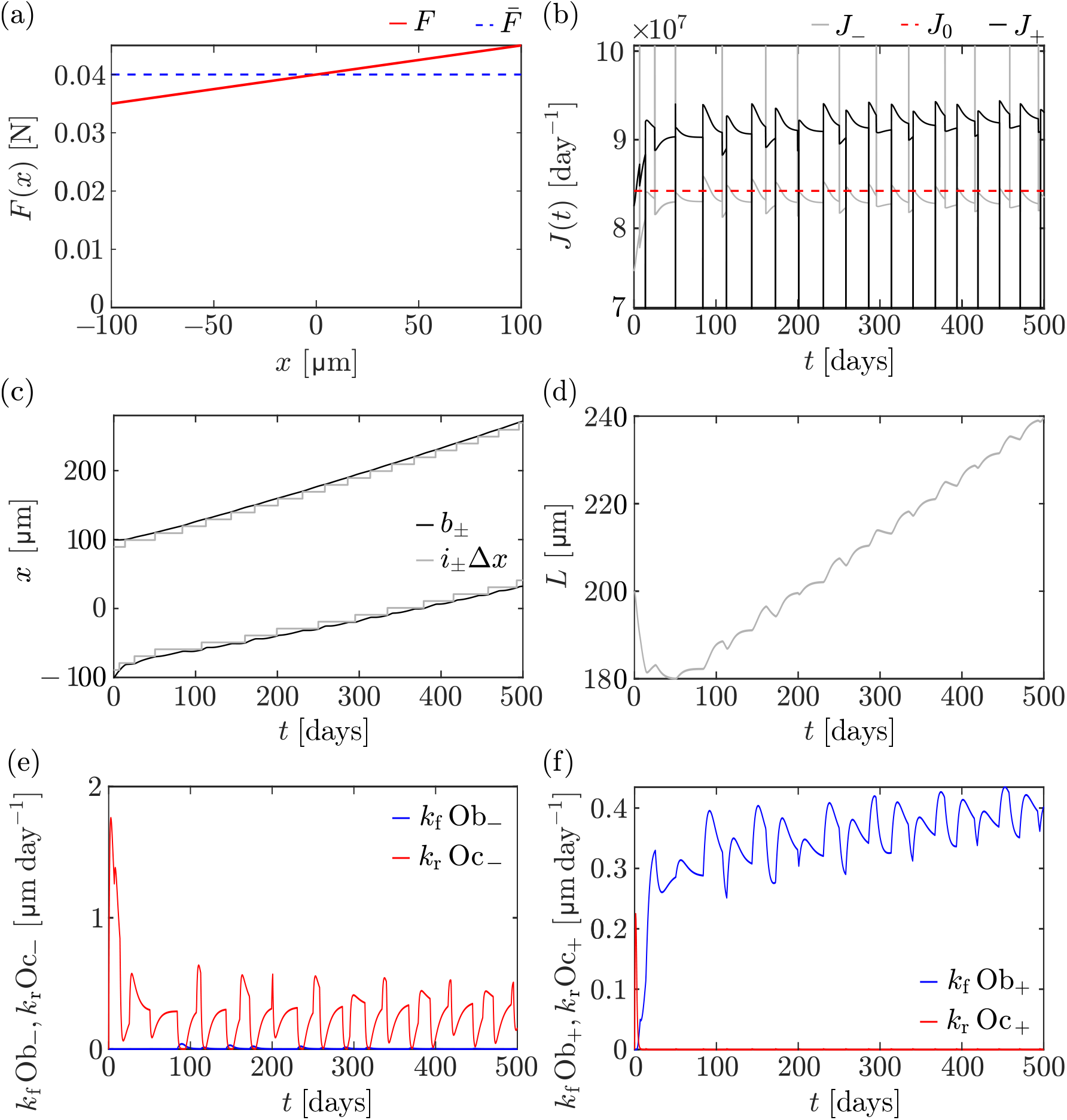
Differential cell generation model with stress stimulation and a force gradient. (a) The applied gradient force (red line) is shown with the reference force 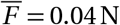 (blue dashed line) over the initial bone domain. (b) Evolution of the fluxes at the left boundary (grey line) and at the right boundary (black line). The reference value *J*_0_ is shown by the red dashed line. (c) Evolution of *i*_±_Δ*x* (grey lines), along with the left and right boundary positions (black lines). (d) Evolution of network length over time. (e–f) Speed of osteoblast-driven (blue line) and osteoclast-driven (red line) boundary movement at the left boundary (e) and right boundary (f) over time

Bone growth during development is known to be significantly stimulated by compressive and bending moments (Carter & Beaupré 2001). The shaft of long bones during growth expands via bone resorption at the endosteal surface and bone apposition at the periosteal surface. Our model reproduces qualitatively these experimental observations that result in an outward drift and thickening of the midshaft bone wall (Carter & Beaupré 2001; Seeman 2008; Maggiano et al. 2015). Our results are also consistent with similar findings from Wolff-law type numerical simulations of the evolution of long bone cross sections under compression and bending (Levenston et al. 1998).

The simulations in Figure 10 suggest that bone in our model does not reach a steady state. The increasing force value with *x* seen by the moving right boundary always induces more bone formation. This is a known shortcoming of commonly used Wolff’s laws and mechanostats, that requires other mechanisms such as osteocyte desensitisation or osteocyte network adaptation to be resolved (Levenston et al. 1998; Turner 1999; Lerebours & Buenzli 2016; Lerebours 2017).

The evolution of the bone boundaries occurs over long timescales, which suggests that the concentration profile of signalling molecules in the network is in a quasi-steady state that evolves slowly due to bone boundary movement. The homeostatic relationship *F*(*L*) in Eq. (33) no longer holds because the force is now inhomogenous, but some insights into the adaptation of bone under a force gradient in our model can be inferred by assuming a quasi-steady state *∂n/∂t* = 0 with the force *F*(*x*) given in Eq. (36). In quasi-steady state, the concentration profile *n*(*x*) of signalling molecules is a solution of Eq. (24) with

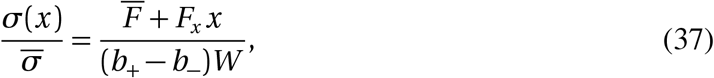

where *b*_+_ and *b*_−_ are assumed to evolve slowly enough to be approximately constant. With the boundary conditions

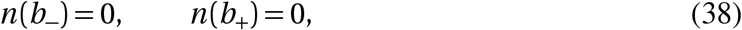

the quasi-steady state solution is given by

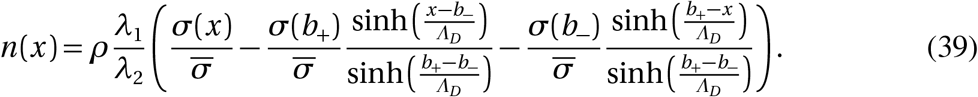

When *b*_−_ = −*L/*2, *b*_+_ = *L/*2 and *F*_*x*_ = 0, Eq. (39) falls back onto Eq. (25). The boundary fluxes *J*_±_ = ∓*D* d*n/*d*x*(*b*_±_) are now functions of both *b*_−_ and *b*_+_ independently, rather than being functions of only the combination *L* = *b*_+_ −*b*_−_, as was the case for a homogeneous loading force:

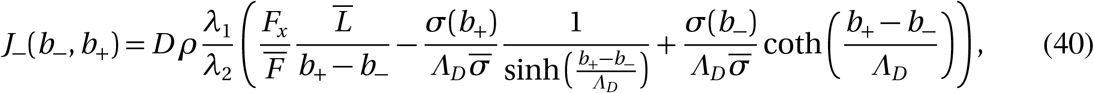

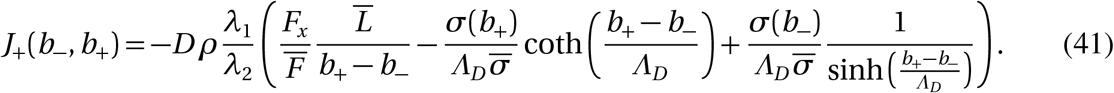

In Figure 11a, we plot the isoflux contour lines *J*_−_ (*b*_−_, *b*_+_) = *J*_0_ and *J*_+_(*b*_−_, *b*_+_) = *J*_0_ in (*b*_−_, *b*_+_) space. A homeostatic steady state is characterised by both boundary fluxes matching the reference value *J*_0_ (see Section 2.3), which is at the intersection of these isoflux contours. The only steady state occurs when *b*_−_ = *b*_+_, i.e., for zero bone length, and at a specific location in space where the force value corresponds to the full-resorption threshold, i.e., where *F*(*b*_−_) = 0.0304 N. Indeed, one can show from Eqs (40)–(41) that

**Figure 11.**
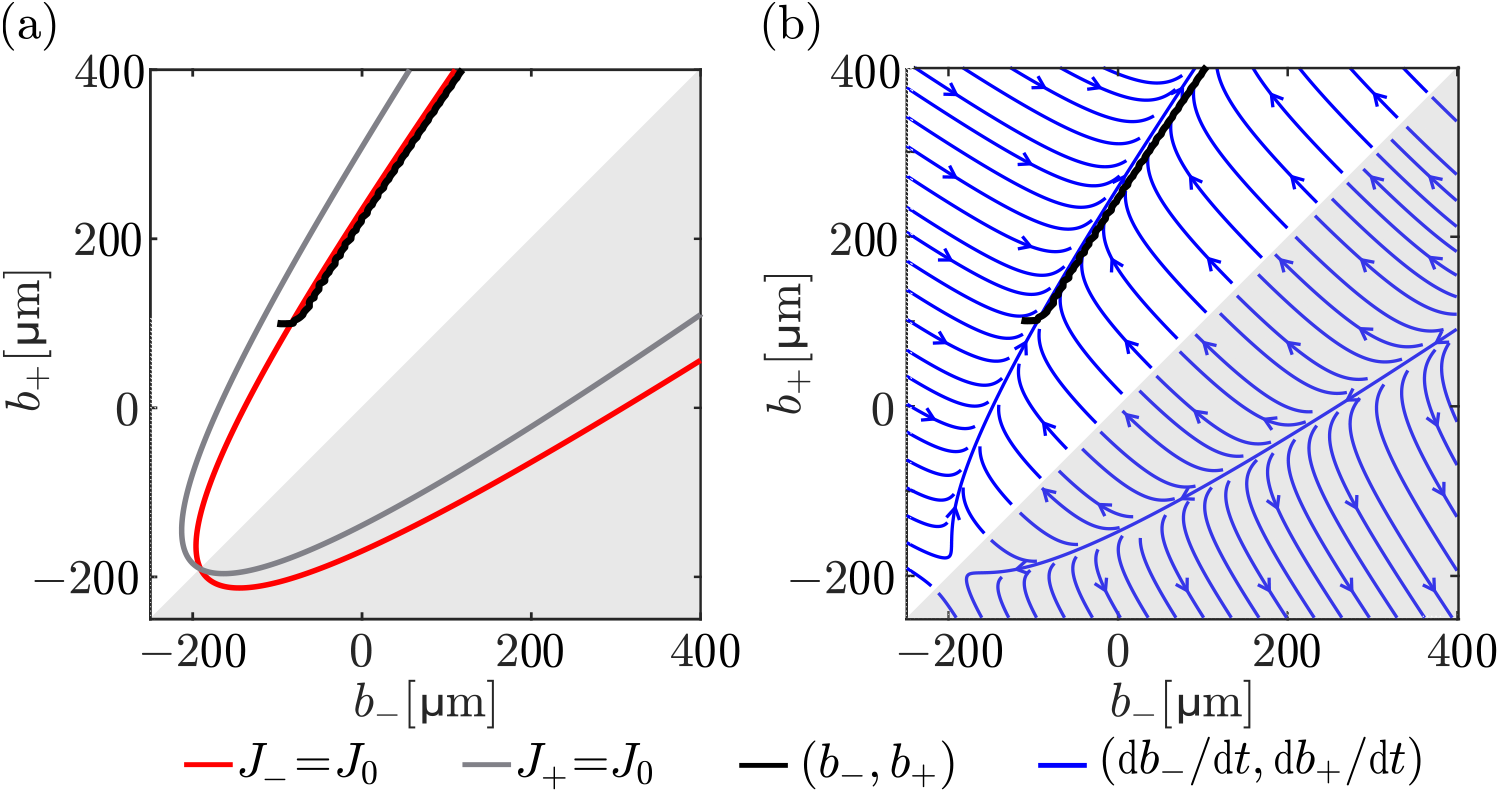
Evolution of boundary positions in t he (*b*_−_, *b*_+_) p lane under a gradient force stimulation. (a) The trajectory of boundary positions (*b*_−_(*t*), *b*_+_(*t*)) corresponding to the simulation in Figure 10 is shown by the solid black line, with contour plots of *J*_−_ = *J*_0_ (red line) and *J*_+_ = *J*_0_ (grey line) obtained from the continuum differential cell generation model in Eqs (40)–(41). These contours intersect at *b*_−_ = *b*_+_ ≈ −190.7µm (see text for more detail); (b) Phase portrait of 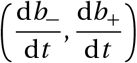 for the instantaneous cell generation model with *v*_*f*_ = *v*_*r*_ = *v* = 10µm/days (blue) and the (*b*_−_ (*t*), *b*_+_(*t*)) trajectory (black) corresponding to the simulation in Figure 10. The grey-shaded areas in (a–b) represent non-physical regions where *b*_−_ *> b*_+_

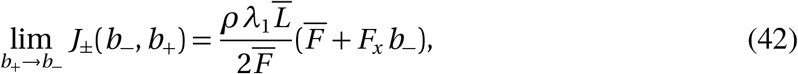

so that when *J*_+_(*b*_−_, *b*_−_) = *J*_−_(*b*_−_, *b*_−_) = *J*_0_ with *J*_0_ = *ρλ*_1_*Λ*_*D*_ tanh 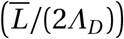 (see Eq. (32)), *b*_−_ is such that

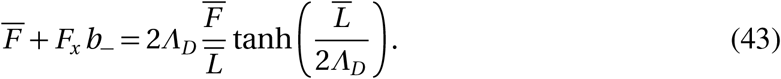

The right hand side in Eq. (43) corresponds to the full-resorption threshold obtained as lim_*L*→0_ *F*(*L*) in Eq. (33). Solving Eq. (43) for *b*_−_ gives the only homeostatic steady state *b*_−_ = *b*_+_ ≈ − 190.7µm.

Figure 11a also depicts the trajectory (*b*_−_(*t*), *b*_+_(*t*)) followed by the numerical simulations of the discrete model (black line). This trajectory closely follows the *J*_−_(*b*_−_, *b*_+_) = *J*_0_ isoflux line, which means that the flux at the left boundary is close to the reference value at all times, as seen also in Figure 10b. However, the position of the left boundary cannot remain stationary. The right boundary flux is always greater than *J*_0_ on average (Figures 10b and 11a). The increase of *b*_+_ due to bone formation at the right boundary forces *b*_−_ to increase as well by following the curve *J*_−_(*b*_−_, *b*_+_) = *J*_0_.

A phase portrait of the evolution of the bone boundaries under the quasi-steady state approximation can be obtained in the continuum limit of the instantaneous cell generation model when *v*_*f*_ = *v*_*r*_ = *v*. In this case, the evolution equations (15)–(17) become

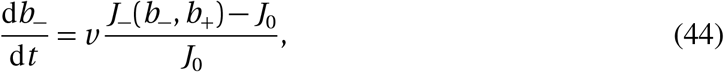

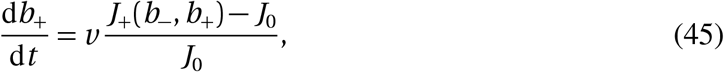

where *J*_±_(*b*_−_, *b*_+_) are given by Eqs (40)–(41). Figure 11b shows that the trajectory *b*_−_ (*t*), *b*_+_(*t*) followed by the discrete simulation with the differential cell generation model follows the field lines of the continuum dynamical system of Eqs (44)–(45) very well. The phase portrait in Figure 11b shows that bone will continue to increase the position of both boundaries while also increasing its length. These results show that while the instantaneous cell generation model leads to instabilities in simulations of the discrete model, its continuum limit may still prove useful in analysing phase portraits of the evolution of bone. Such analyses may provide good indications of how further mechanisms may ensure that bone does not grow unbounded under compression and bending.

## 4 Conclusion

This paper presents a novel mathematical model of bone adaptation that explicitly takes into account the propagation of signalling molecules in the osteocyte network as a mechanism to induce bone formation or bone resorption. The model is based on a one-dimensional osteocyte network, where molecules are emitted by osteocytes in response to mechanical stimulus, and where these molecules jump from one osteocyte to the next until they reach the bone surface. The explicit inclusion of signal propagation through a dynamic network of osteocytes in this model leads to new behaviours compared to previous continuum models of bone adaptation. Cycles of unloading and reloading may not always restore bone to its initial state. Furthermore, when osteocytes are stimulated by mechanical stress, our model predicts the emergence of a minimum threshold of loading force below which all bone is resorbed. However, no such minimum threshold is predicted when osteocytes are stimulated by the strain energy density. These kinds of qualitatively distinct behaviours may be useful in determining what mechanical cues osteocytes are most sensitive to in responding to mechanical stimulus (Chen et al. 2010). These cues still remain largely unknown, particularly for networks composed of few osteocytes, such as in trabecular struts.

While our model does not currently consider physiological disruptions that occur during age-related bone loss, it would be interesting to extend our model to investigate the mechanical adaptation of bone during endocortical bone loss. Mechanical feedback following age-related bone loss at the endosteal surface is believed to induce bone apposition at the periosteal surface, resulting in a similar drift of the cortical wall obtained in our model under a force gradient, except with a thinning of the bone wall (Stein et al. 1998; Seeman 2008; Boskey & Coleman 2010; Buenzli et al. 2013). Such extensions of our model would provide insights into the role of osteocyte disruptions during this mechanical adaptation, such as decreases in osteocyte numbers observed experimentally (Jilka, Noble & Weinstein 2013).

The consideration of osteocyte creation during bone formation and osteocyte removal during bone resorption leads to regular disruptions in signalling molecules emanating from the bone surface, which in turn leads to periodic variations in the populations of osteoblasts and osteoclasts. Much remains to be understood about mechanisms that lead to the formation of lamellae in lamellar bone, but it possible that lamella formation may be linked to behaviours enabled by the discrete nature of osteocytes and their effect on signal propagation.

Our mathematical model has a number of limitations. First, it only considers the evolution of bone length and the propagation of signals in a one-dimensional network. Osteocytes are also assumed to be regularly spaced. While there is some evidence that osteocytes are arranged in layers (Marotti 2000; Ardizzoni 2001), it is not clear what mechanisms may lead to such an arrangement. In our model, we assumed that osteocytes may be stimulated mechanically either by mechanical stress, or by strain energy density, as has been commonly assumed in several works (Cheong et al. 2021; Smotrova & Silberschmidt 2022, 2024). Other works consider other types of mechanical stimulations, such as fluid-flow induced shear stress and dependences upon strain rates (Carter & Beaupré 2001; McGarry et al. 2005; Adachi et al. 2010; van Tol et al. 2020a,b). It is likely that a variety of mechanical stimulations may induce bone formation and bone resorption in reality (Chen et al. 2010). The consideration of other types of mechanical stimulations in our model could lead to yet other behaviours of bone adaptation than those explored in our work. Finally, our model suffers from similar shortcomings as standard mechanostat theories and Wolff’s laws, in which full bone resorption occurs under complete disuse, and unbounded bone growth occurs under bending (Levenston et al. 1998; Turner 1999; Skerry 2008; Lerebours 2017). To address these shortcomings, future works may need to account for osteocyte desensitisation and osteocyte network adaptation (Turner 1999; Lerebours & Buenzli 2016; Pauchard & Buenzli 2025).

Our model focusses on the propagation of intercellular signals associated with mechano-response, rather than the transport of fluid involved in mechano-sensation. Future avenues for this model include considering the coupling of these two network transport mechanisms, i.e., mechano-sensation by fluid flow, and mechano-response through the cellular network, because they both depend on the architecture of the lacuno-canalicular and osteocyte networks. Efficient mathematical models of fluid flow within the LCN currently rely on Kirchhoff’s circuit theory to represent conservation of current at branching points of the network (van Tol et al. 2020a,b; Fu & Yang 2024). Our mathematical model could be adapted to represent fluid flow based on biased jump rates of fluid particles or of response-triggering molecules between osteocyte lacunae. Such models of propagation through networks are similarly conservative, and can lead to a reaction–diffusion–advection equation in an appropriate continuum limit (Mehrpooya, Challis & Buenzli 2024), where advection may represent fluid flow.

In conclusion, our mathematical model provides a first model of the regulatory control of the osteocyte network for bone adaptation to mechanical loads. This model illustrates of the importance of accounting for the discrete nature of osteocytes during bone formation and bone resorption.

## Acknowledgements

We thank Richard Weinkamer and Nathalie Bock for fruitful discussions. PRB acknowledges support from the Australian Research Council (DP190102545). AM acknowledges support by a QUT Postgraduate Research Award. AM, VJC, and PRB acknowledge support from the Max Planck Queensland Centre on the Materials Science of Extracellular Matrices (MPQC).

## Appendix A Osteoblast and osteoclast parameter estimation

### A.1 Instantaneous cell generation model

We examine the instantaneous cell generation model when the loading force is increased by 20% at *t* = 40 days, from *F* = 0.04 N to *F* = 0.048 N. For illustration, we assume equal formation velocity *v*_f_ and resorption velocity *v*_r_. Figure 12a shows the evolution the right boundary position *b*_+_(*t*) over a short period of time following the overload. The force increase at *t* = 40 days leads to a right boundary flux *J*_+_(*t*) *> J*_0_ (Figure 12b) which induces bone formation (Figure 12a). When sufficient bone is formed to create a new osteocyte, this new osteocyte has no signalling molecules initially. The flux *J*_+_ drops to zero (Figure 12b). This immediately leads to bone resorption in the instantaneous cell generation model by Eq. (17), which in turn triggers the resorption of the newly formed osteocyte. Since that osteocyte has collected signalling molecules meanwhile, its resorption frees extra signalling molecules and the flux increases again above *J*_0_, leading to bone formation. The process repeats and leads to unstable oscillations in both the boundary position *b*_+_(*t*) and the boundary flux *J*_+_(*t*). Our investigations showed that this behaviour cannot be remedied by choosing different values of *v*_f_ and *v*_r_. Such fast responses of the model to changes in *J*_+_(*t*) are biologically unreasonable, since populations of osteoblasts and osteoclasts take time to generate and to eliminate.

**Figure 12.**
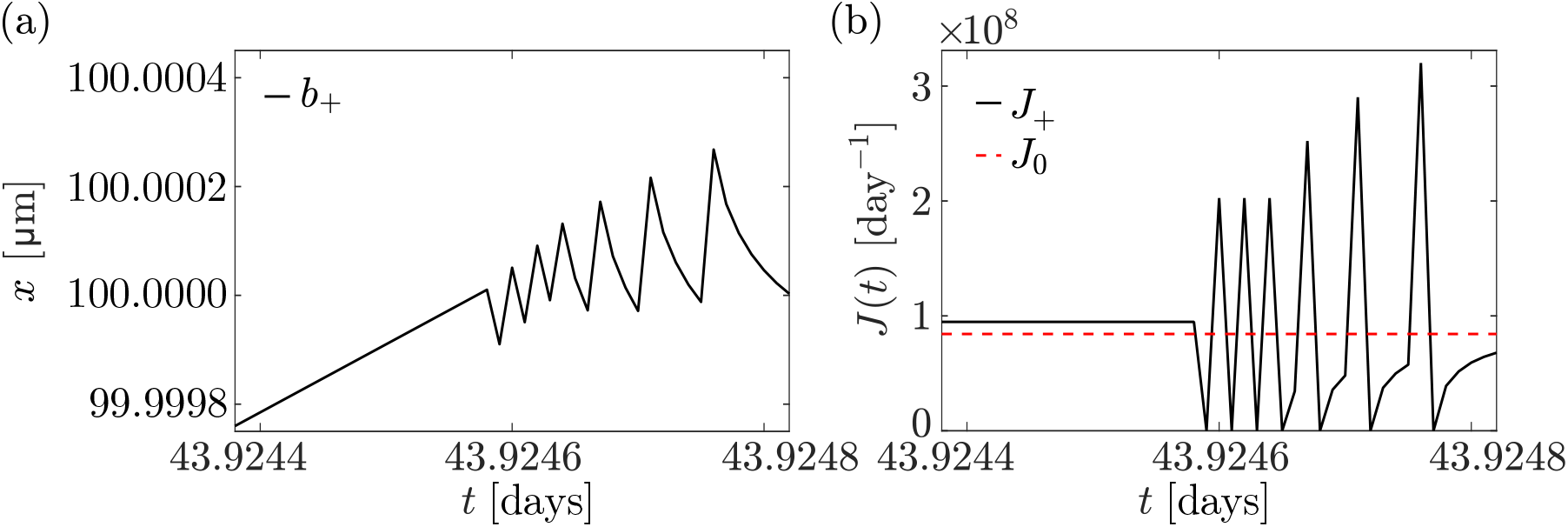
Overloading the instantaneous cell generation model with a 20 % force increase from *F* = 0.04 N to *F* = 0.048 N at *t* = 40 days. (a) Evolution of the right boundary position (black line) over time. (b) Evolution of the right boundary flux (black line) over time corresponding to (a), with the reference value *J*_0_ shown (red dashed line). Parameters (in addition to Table 1): *v*_f_ = *v*_r_ = 10µm/days

### A.2 Differential cell generation model

Compared to the instantaneous cell generation model, the differential cell generation model provides a more controlled response to overload by regulating boundary motion through the gradual accumulation or elimination of osteoblasts and osteoclasts.

Oscillations in boundary positions may still occur in the differential cell generation model. However, unlike in the instantaneous cell generation model, these oscillations remain stable and diminish with time. Figure 13 presents the evolution of the right boundary *b*_+_(*t*), flux *J*_+_(*t*), and cell populations Ob(*t*) and Oc(*t*) when the loading force is increased by 20% from *F* = 0.04 N to *F* = 0.048 N at *t* = 40 days. Two different parameter regimes are shown. Figure 13a corresponds to an underdamped regime where cell elimination is weak. The boundary flux oscillates, leading to oscillations in the boundary position. Figure 13b corresponds to an overdamped regime, where cell elimination is strong. In this regime, there are no oscillations in the boundary position.

**Figure 13.**
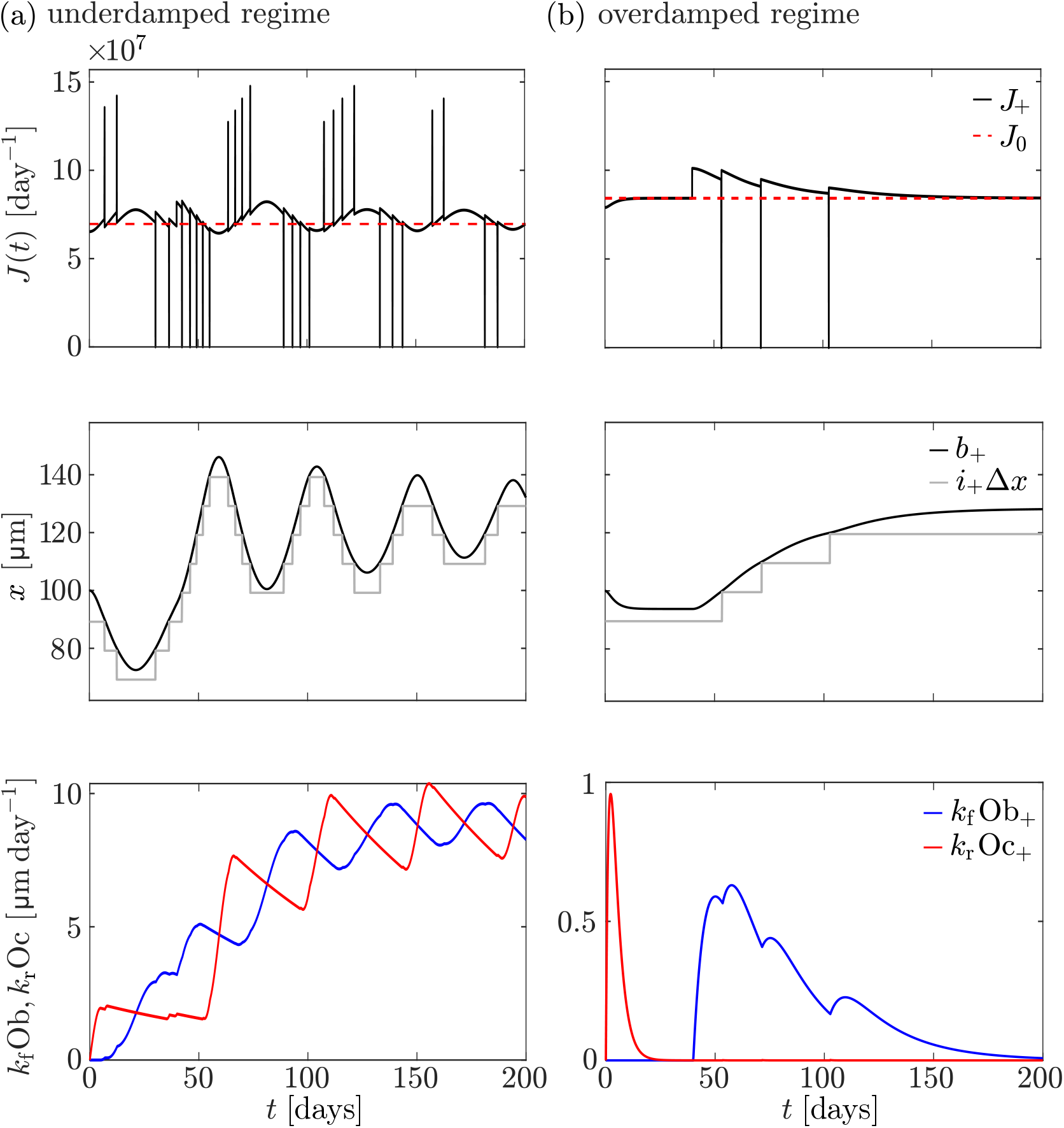
Overloading the differential cell generation model with a 20 % force increase from *F* = 0.04 N to *F* = 0.048 N at *t* = 40 days. Each column includes plots of evolution of the right boundary flux (black line) over time with the reference value *J*_0_ shown (red dashed line), evolution of *i*_+_Δ*x* (grey line) and the right boundary position (black line) over time, and speed of osteoblast-driven (blue line) and osteoclast-driven (red line) boundary movement at the right boundary over time. (a) Underdamped regime with *A*_Ob_ = *A*_Oc_ = 0.01 /days. (b) Overdamped regime with *A*_Ob_ = *A*_Oc_ = 0.5 /days. All other parameters are as per Table 1

Some insights into when the parameters are in an underdamped regime and when they are in an overdamped regime can be gained when the parameters satisfy some symmetry conditions, as detailed in Section A.2.1.

#### A.2.1 Criterion for overdamped regime

The numerical simulations presented in Figure 13 suggest that the cell elimination rate in Eqs. (20) and (21) helps control oscillations in the boundary position and flux. To understand why, we seek a mathematical expression that characterises the overdamped regime under further assumptions on parameter values in which bone formation and bone resorption parameters have similar values. While this is not what is assumed in simulations presented in our paper, the analysis made under these assumptions still provides a good indication of how parameters affect oscillation damping.

Since the loading force and initial condition are symmetric, Ob_+_(*t*) = Ob_−_(*t*) and Oc_+_(*t*) = Oc_−_(*t*). The evolution of network length over time is governed by:

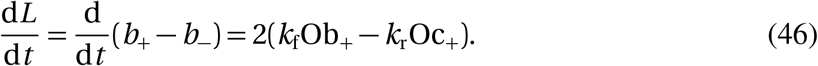

Differentiating with respect to time and using Eqs (18)–(19), we have

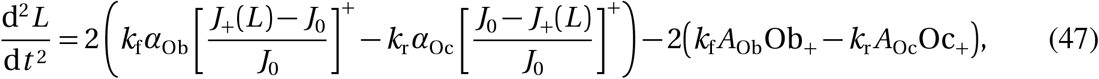

where we wrote the dependence of *J*_+_ upon *L* explicitly. To simplify the analysis of the behaviour of Eq. (47), we now assume that

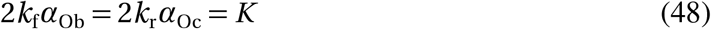

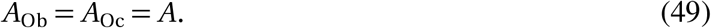

Since [*x*]^+^ − [−*x*]^+^ = *x* for any value *x*, Eq. (47) becomes

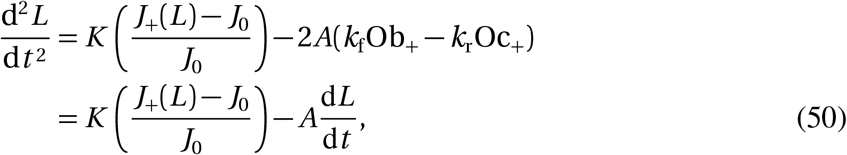

where we used Eq. (46) for the second equality. Equation (50) corresponds to the evolution equation of a nonlinear damped oscillator. Its oscillatory behaviour around the homeostatic steady state 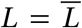 when 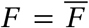 can be assessed by linearising *J*_+_(*L*) about 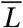. For this, we assume that *J*_+_(*L*) can be approximated by the steady-state expression in Eq. (27) (since molecule propagation dynamics is much faster than bone length changes). With Eq. (32), we can write, when 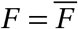,

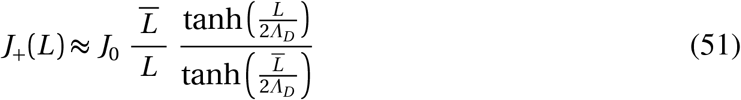

Linearising Eq. (51) about 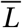 gives

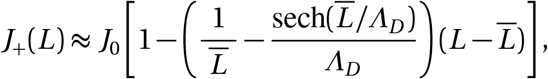

so that linearising Eq. (50) about 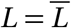 gives

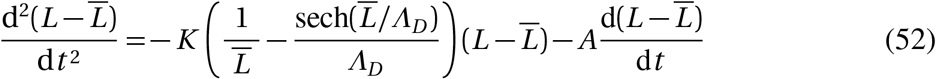

Equation (52) is the evolution equation of a damped harmonic oscillator for the deviation 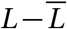. Its overdamped regime is obtained when

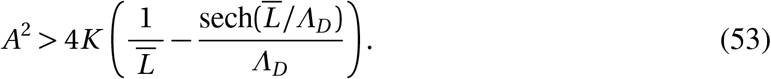

